# Plasticity in thoracic paravertebral sympathetic postganglionic neurons after high spinal cord transection

**DOI:** 10.1101/2025.09.06.674676

**Authors:** Yaqing Li, Krishna Pusuluri, Mallika Halder, Alan Sokoloff, Astrid A Prinz, Shawn Hochman

## Abstract

Various pre-sympathetic descending brain circuits recruit spinal cord preganglionic neurons to encode central sympathetic drive via their synaptic actions onto sympathetic postganglionic neurons (SPNs) - the final sympathetic output neurons. Thoracic paravertebral ganglia SPNs (tSPNs) provide distributed control over body tissue systems via functional subpopulations. High thoracic spinal cord injuries (SCIs) compromise descending excitatory drive to SPNs, causing dysautonomias including hypotension. In adult mice, we tested whether the SCI-induced chronic reduction in tSPN activity leads to homeostatic increases in their excitability. tSPN excitability spanned a >10 fold range in both sham and SCI populations, governed by a strong linear (ohmic) relationship between cell resistance and threshold depolarizing current (rheobase), with a clear trend towards increased excitability after SCI. Dendritic length was reduced, as was measured cell capacitance in Neuropeptide Y expressing (NPY^+^) tSPNs (putative vasoconstrictors), which represent >40% of tSPNs. NPY^+^ tSPNs also had changes in active membrane properties including an increased repetitive firing output gain (↑*f*-I slope), which modelling attributed to reduced delayed rectifier currents (I_K_). After SCI, spontaneous quantal excitatory synaptic frequency increased overall (226%) and in the NPY^+^ tSPN subpopulation (300%); their temporal summation recruited spiking in 10.5% of sham and 22.2% of SCI recordings. Computational modeling showed that spontaneous synaptic activity was particularly effective at recruiting spiking after SCI. Overall, tSPNs, including in vasoconstrictors, appear to undergo CNS-independent compensatory increases in excitability after SCI. The alterations further contribute to observed central and peripheral changes that limit hypoactivity and hypotension but exaggerate reflex responses.

**Significance Statement:** Recruited by preganglionic neurons located in thoracolumbar spinal cord, sympathetic postganglionic neurons (SPNs) represent the final step in sympathetic neural homeostatic control of target organs. The dramatic reductions or complete loss of brain pre-sympathetic drive in higher level spinal cord injuries (SCIs) contribute to emergent dysautonomias. Given the crucial role of SPNs in maintaining organismal homeostasis, we undertook comprehensive studies to test whether thoracic SPNs undergo homeostatic compensatory increases in their cellular excitability weeks after a high thoracic SCI. We observed changes consistent with increased cellular and synaptic excitability including in SPN vasoconstrictors, whose increased output gain, could mitigate hypotension but also strengthen hypertensive responses generated by afferent-driven exaggerated preganglionic drive (e.g. autonomic dysreflexia).

## Introduction

The final output of the central sympathetic system originates from spinal preganglionic neurons located in the thoracic and upper lumbar spinal cord. These spinal sympathetic preganglionic neurons project to sympathetic postganglionic neurons (SPNs) in paravertebral and prevertebral ganglia, which in turn innervate various organ systems. In individuals with cervical or high thoracic spinal cord injury (SCI), there is a partial or complete loss of supraspinal control to the spinal sympathetic neural output, leading to dysfunction in multiple organ systems, including, thermoregulatory, digestive and cardiovascular systems^1–4^. Vasomotor dysregulation in SCI individuals is characterized by persistent orthostatic hypotension and episodic hypertensive crises known as autonomic dysreflexia (AD). AD is a potential life-threatening hypertensive event mediated predominantly by visceral nociceptive-fiber-induced bouts of uncontrolled excessive sympathetic activities^5,6^. Studies on the mechanisms underlying SCI-induced exaggerated vasomotor function have identified plasticity in spinal cord afferent projections and circuits that act on spinal sympathetic preganglionic neurons^7–9^. Additional mechanisms in the periphery that strengthen vasomotor tone include reduced noradrenaline reuptake and increased adrenergic receptor sensitivities at arteries^10,11^, and an exaggerated blood pressure control through the renal angiotensin system^12^. Notably absent from these studies, however, are considerations of changes in the excitability of SPNs, the final arbiter of neural actions on effector tissues.

Paravertebral thoracic sympathetic postganglionic neurons (tSPNs) prominently innervate widely distributed organ systems, including the vasculature in skeletal muscle and skin^13^. While spinal preganglionic axons diverge to innervate many tSPNs, the only presumed function of the numerically greater tSPNs (14:1 in mouse; 200:1 in human) is to faithfully amplify and spatially spread activity along target organs. Even though paravertebral SPNs receive convergent inputs from multiple preganglionic neurons^14^, conventional dogma is that one *primary* synapse is thought to provide a *supra*threshold excitatory postsynaptic potential (EPSP), with *secondary* synapses generating wholly inconsequential subthreshold EPSPs^15^. Whether mammalian synaptic organization adheres to the dogma of primary and secondary synapses in all tSPNs is not obvious^16–18^. Indeed, incorporation of more accurate measures of membrane properties using whole-cell recordings support a much greater capacity for synaptic integration in output gain modulation than previously considered^19,20 21,22^. Observations of SPNs as not merely followers but as dynamic contributors to the regulation of sympathetic output to peripheral organs have also been seen^17,23^. Overall, it is important to determine whether changes in tSPN excitability after high thoracic SCI contribute to sympathetic dysfunction.

The present study undertook high thoracic spinal transection in adult mice, a SCI model known to induce various dysautonomias including hypotension and AD ^24–26^. Using an *ex vivo* approach and focusing on whole-cell patch recordings in individual tSPN neurons, we undertook a comprehensive assessment of their intrinsic excitability changes after SCI. We also assessed their morphological plasticity in tyrosine hydroxylase (TH)^TdTomato^ mice with sparse labeling^27^.

Recognizing that tSPNs can be subdivided into 7 distinct subpopulations and that the small-diameter neuropeptide Y (NPY)-expressing SPNs predominantly innervate vasculature as vasoconstrictors ^28–30^, we further specifically explored injury-induced plasticity in NPY^+^ tSPNs. Additional recordings from rostral and caudal thoracic ganglia in the intact thoracic chain investigated population SPN responses. An optimized computational model of tSPNs^19^ was used to further explore possible injury-induced intrinsic and synaptic changes. Overall results demonstrate that tSPNs are highly heterogeneous and, while surprisingly resilient to SCI, undergo changes consistent with homeostatic compensatory increases in excitability.

## Methods

### Animals

All animal procedures were approved by the Emory University Institutional Animal Care and Use Committee in accordance with regulations of the Guide for the Care and Use of Laboratory Animals. Experiments were conducted using adult C57BL/6 mice (P80-156), encompassing both male and female mice in all experimental groups. The gender distribution was approximately 1:1 across all examined groups. NPY^TdTomato^ mice were used for NPY^+^ tSPNs whole-cell recording, by crossing NPY^Cre^ mice (Jax# 027851) with R26^TdTomato^ mice (Jax# 007908). Reconstruction of adrenergic tSPNs was accomplished in TH^TdTomato^ by crossing sparse-labelling TH^Cre^ mice (Jax# 008601) with R26^TdTomato^ mice (Jax# 007908). Choline acetyltransferase (ChAT)^ChR^ mice were utilized for tSPNs compound potential recordings, by crossing ChAT^Cre^ mice (Jax# 006410) with R26^ChR2-eYFP^ mice (Jax# 024109).

### Spinal cord injury

Mice (∼P80-100) were anesthetized using inhaled isoflurane. Under sterile conditions, the T2 vertebrae was localized by touch, followed by vertebrae laminectomy to expose the T2-3 spinal cord. For the spinal cord injury group, the T2 spinal cord was completely severed with sterile iridectomy scissors to create a complete transection. In the sham group, vertebrae were removed without spinal cord transection using littermates of the SCI mice. After the surgical procedure, the overlying muscle was moved back to the original location, and the skin wound was sutured by VetBound. Both groups of animals received analgesic (buprenorphine, 0.05mg/Kg, I.P.) mixed with body temperature sterile lactated Ringer Solution to prevent dehydration (0.5ml) immediately after surgery. The animals were then allowed to recover on a temperature-controlled heating pad, and analgesic and Ringer Solution were administered for two days with 12 hours apart. Later, Ringer Solution was continuously given to SCI mice twice a day for at least 2 weeks to prevent dehydration. Daily inspection and manual bladder expression were performed twice a day until the terminal experiments.

### Electrophysiology

Experiments were conducted at 3wks, 6wks or longer after SCI surgery or 3 to 6 weeks after sham surgery. Age-matched un-operated mice were used as the naïve group. Notably, no significant differences were observed between the early (3wks post SCI) and late (6-8wks post SCI) chronic SCI groups (not shown), so these data were pooled together and are represented as the SCI group.

#### Tissue Preparation

Mice were initially anesthetized with inhaled isoflurane and subsequently maintained under urethane anesthesia (i.p. injection, 2000mg/kg). Complete sedation was confirmed by the absence of foot pinch and eye blink reflex. The thoracic spinal column, along with all ribs, were quickly dissected out and placed in a dissection chamber filled with continuously oxygenated dissection solution. **Figure 1A** presents a simplified schematic of the anatomic organization of intraspinal preganglionic and paravertebral postganglionic neurons. The spinal cord was removed from the spinal column following a dorsal laminectomy. Then the ventral spinal column was cut along the midline after the removal of aorta. This halved vertebral column attached to the ipsi-side ribs was kept for experiments to maintain the integrity of the thoracic paravertebral chain.

**Figure 1.**
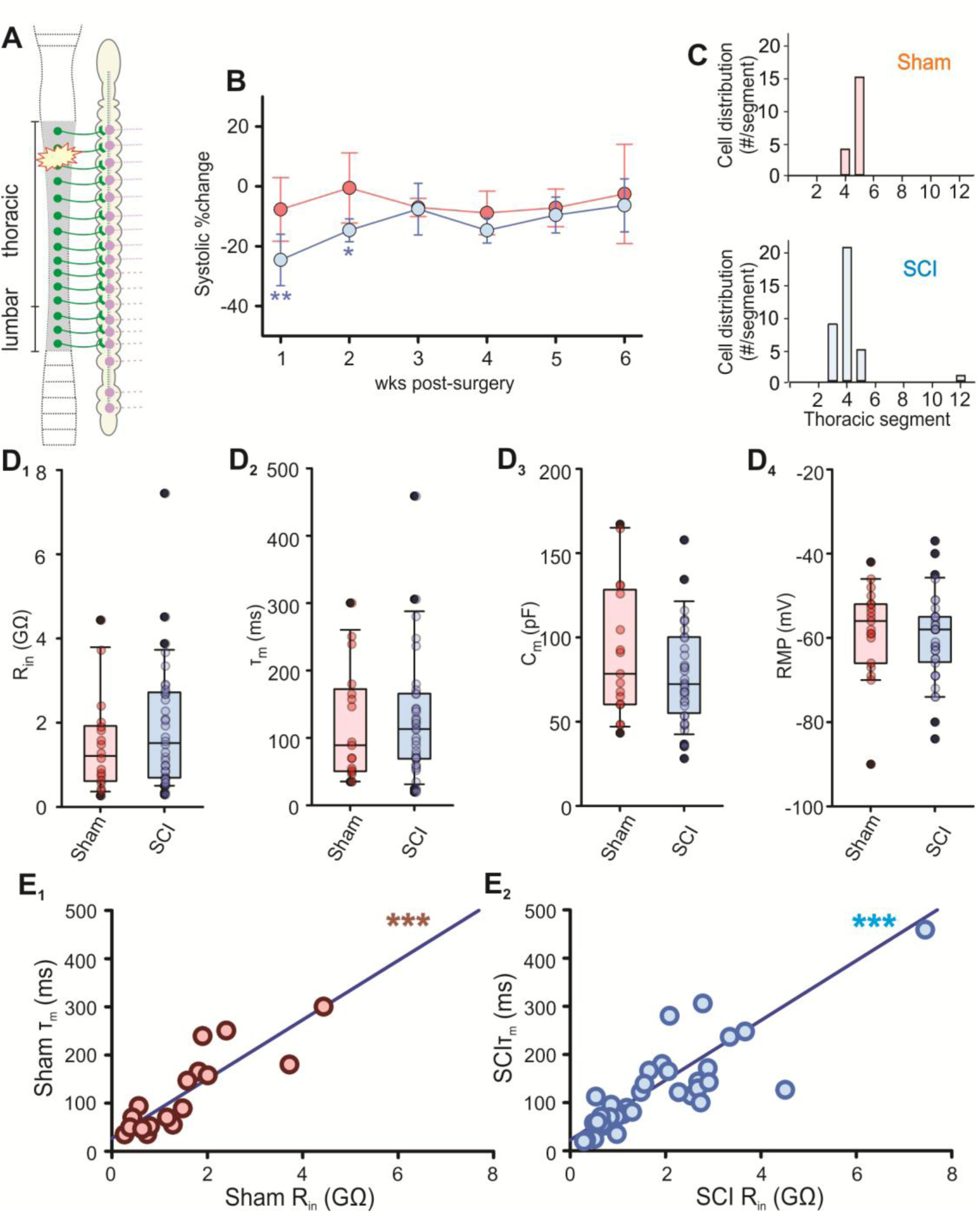
Whole population passive properties. The whole population tSPN passive properties in sham and chronic spinal cord injury groups. ***A,*** Schematic of the thoracic paravertebral chain and its received central input. SPNs (purple) in the thoracic paravertebral chain receive central input from spinal sympathetic preganglionic neurons (green). ***B,*** Temporal change of systolic pressure in sham (red, n = 8) and SCI (blue, n = 16) mice post-surgery. Systolic pressure was normalized with pre-surgery pressure. SCI mice had a significantly reduced systolic pressure in the first two weeks after SCI (P = 0.002 at 1wk po-SCI and P = 0.013 at 2wk po-SCI). * Indicates significance compared to baseline. Unpaired t-test. ***C,*** Histogram showing the number of whole-cell recorded tSPNs in each thoracic ganglion in sham (n = 6 mice) and chronic SCI (n = 15 mice) groups. ***D,*** Summary of passive properties of whole population tSPNs in sham and SCI groups. No significant difference was observed in cellular input resistance (***D_1_***, R_in_, P = 0.30), time constant (***D_2_***, τ_m_, P = 0.62), capacitance (***D_3_***, C_m_, P = 0.44) and resting membrane potential (***D_4_***, RMP, P = 0.53). Unpaired student t-test. Box plots and horizontal bar within represent the interquartile range and median, respectively. Error bars extend to the most extreme data point that is within 1.5 times the interquartile range. ***E,*** R_in_ was strongly correlated with τ_m_ in both sham (***E_1_***, r = 0.86, P = 1.1×10^-5^) and SCI (***E_2_***, r = 0.76, P = 9.6×10^-8^) groups. Solid line indicates linear simple regression. Pearson correlation test. * P < 0.05, ** P < 0.01, *** P < 0.001.

**Figure 2.**
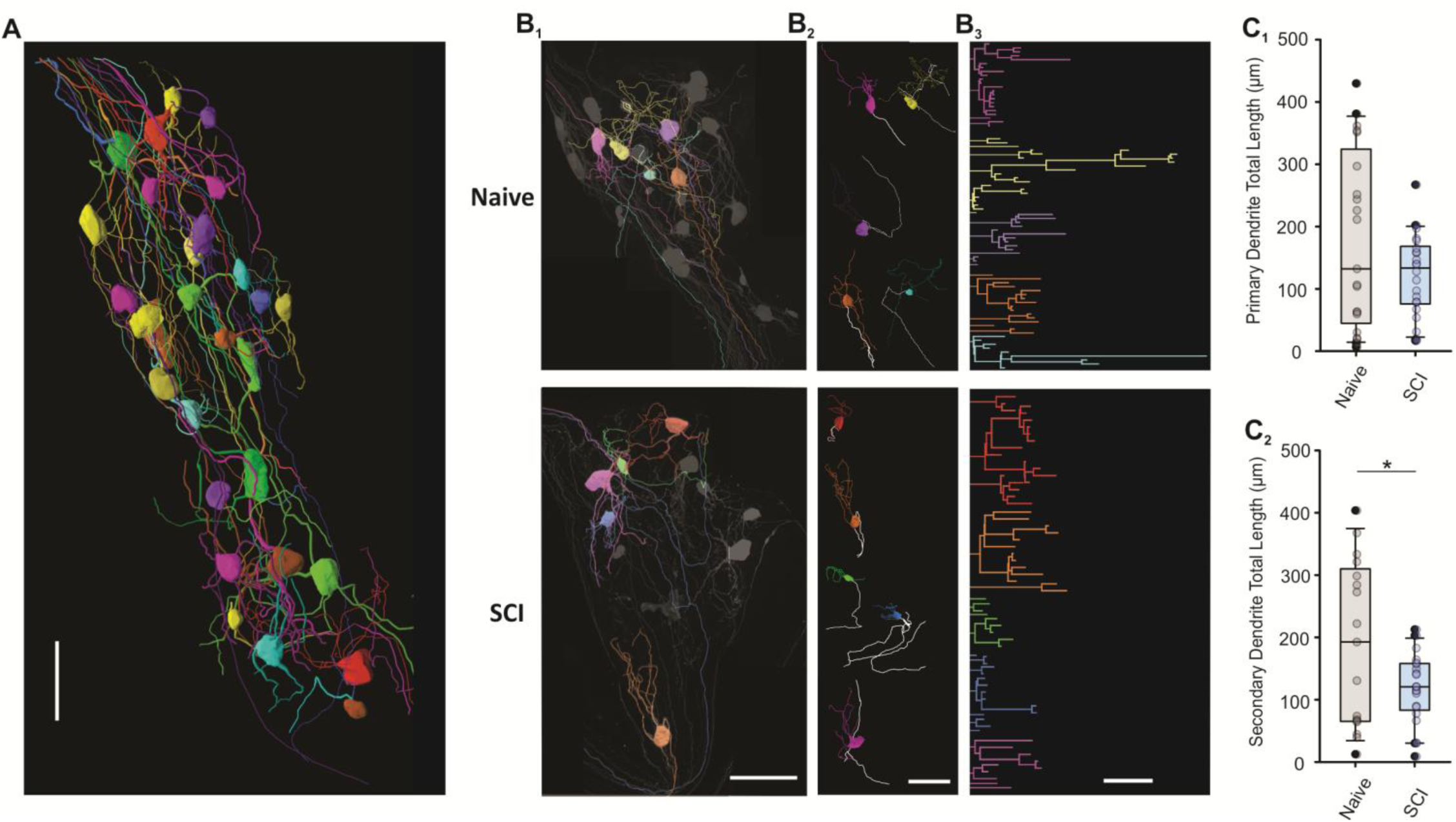
Anatomy-3D reconstruction. 3D-reconstruction of TH^+^ tSPNs in naive (n = 3 mice) and 3 weeks post-SCI (n = 4 mice) group. ***A,*** Example of 3D-reconstruction of TH^+^ tSPNs from a naive mouse. Each ganglion had a limited number of tSPNs labeled, showing a large variability in soma size, neurite number and length. Scale bar 50μm. ***B,*** Examples of dendrite reconstruction. ***B_1_,*** Examples of sparse labeled TH^+^ tSPNs (gray) from a T4 ganglion in a naive mouse and a T4 ganglion in SCI mouse. Five tSPNs (colored) from each ganglion were randomly selected from labeled tSPNs as examples to show detailed neurites reconstruction in ***B_2_,*** where dendrites were colored, and axons were marked white. ***B_3_,*** Schematic reconstruction of colored tSPNs dendrites from B_2_. Dendrite diameter not to scale. Scale bar 100μm. ***C,*** Summary of dendrite length from reconstructed TH^+^ tSPNs. ***C_1_,*** No significant difference in primary dendrite length between naive and SCI, P = 0.112. ***C_2_,*** Significant reduction in secondary dendrite length after SCI, P = 0.035, unpaired student t-test. * P < 0.05

#### Whole-cell recordings

The dissected tissue was incubated in continuously oxygenated HEPES solution containing collagenase (type III, 20mg per 1mL aCSF, Worthington Biochemical Corporation) at 37C° for 1 hour. Following incubation, the tissue was vortexed to remove adherent fat and washed with dissection solution several times to eliminate residual collagenase. The intact sympathetic chain was then separated by severing rami in dissection solution, and then transferred and pinned down into a clear Sylgard recording dish. Recirculating, oxygenated recording artificial cerebrospinal fluid (aCSF) was continuously perfused through the dish and tissue was allowed to rest at least 30mins before recording.

Whole-cell patch recordings were conducted from postganglionic neurons at room temperature. Identification of neurons was performed using an upright microscope (Olympus, BX51WI) equipped with a low-light camera (Olympus, OLY-150). Patch electrodes were pulled using a vertical puller (Narishige, PP-83) from 1.5mm outer diameter filamented, borosilicate glass capillaries (World Precision Instruments, stock # TW150F-4) to achieve a target resistance of 5-9 MΩ with electrode solution. Signals were amplified using a MultiClamp 700A and digitized at 10 kHz using a Digidata 1322A and Clampex software (Molecular Devices).

For analysis, we considered all neurons displaying clearly defined action potentials upon depolarization by square current steps. Of these, cells were excluded 1) if more than 100pA was required to hyperpolarize a cell to −70mV (indicative of a significant leak), 2) if action potentials appeared stunted (indicative of an incomplete breakthrough), or 3) if membrane potential was highly variable (indicative of improper seal formation). All included neurons meeting these criteria exhibited input resistances higher than 250MΩ.

Recordings for the whole population of tSPNs were obtained from 19 neurons in 6 sham mice and 36 neurons in 15 SCI mice. NPY^+^ tSPN recordings were obtained in 12 neurons in 6 sham mice and 12 neurons in 8 SCI mice. Due to technical challenges in these experiments, we opted to include recordings from NPY^+^ tSPNs in the whole population tSPN recordings. According to a prior RNAseq study, approximately 50% of tSPNs express NPY ^28^. Therefore, in our experiments using NPY^TdTomato^ mice to target NPY^+^ tSPNs, to prevent bias towards the NPY^+^ population, we recorded an equivalent number of NPY^-^ tSPNs from each animal. This approach allowed us to effectively integrate the recordings from NPY^+^ tSPNs into the overall whole population statistical analysis.

#### Population recordings

Briefly, the right side of the thoracic sympathetic chain with the right rib cage was transferred to oxygenated recording aCSF at room temperature. The interganglionic nerve was carefully dissected and cut either rostral to T4 ganglion, or caudal to T12 ganglion, and allow the tissue to rest for 30mins. A glass suction electrode was filled with recording aCSF and placed with a mild suction on the interganglionic nerve either towards the T4 or T12 ganglion for recording. Evoked population T10 spinal preganglionic responses were recorded following supramaximal electrical stimulation of their axons in the T10 ventral root. Compound action potential (CAP) response magnitudes were determined following signal rectification and integration and subtraction of baseline noise (subtracting an equal duration of rectified and integrated baseline voltage fluctuations). Recorded CAP volleys comprised both direct recruitment of rostrally (T4/5) and caudally (T11/12) projecting preganglionic axons and synaptic recruitment of adjacent ganglionic tSPNs. The preganglionic component of CAPs was obtained after block of synaptic transmission with hexamethonium (Hex; 50-100 μM) while the synaptically-recruited tSPN recruitment was calculated as the difference in CAP amplitude before and after Hex. Evoked responses were captured every 60 seconds and the average response of 10 episodes was compared before and after Hex.

#### Solutions

Dissection solution contains [in mM]: NaCl [128], KCl [1.9], MgSO_4_ ·7H_2_O [6.5], CaCl_2_·2H_2_O [0.84], KH_2_PO_4_ [1.2], glucose [10], and NaHCO_3_ [26]. Incubation HEPES solution contains [in mM]: NaCl [92], KCl [2.5], NaH_2_PO_4_ [1.2], MgSO_4_ ·7H_2_O [2], CaCl_2_·2H_2_O [2], NaHCO_3_ [30], HEPES [20], glucose [25], sodium ascorbate [5], thiourea [2], sodium pyruvate [3]. All recordings were made in recording aCSF containing [in mM]: NaCl [128], KCl [1.9], MgSO_4_ ·7H_2_O [1.3], CaCl_2_·2H_2_O [2.4], KH_2_PO_4_ [1.2], glucose [10], and NaHCO_3_ [26]. pH of above three solutions was adjusted to 7.4 after saturation with gas (95%O_2_, 5%CO_2_) at room temperature. Intracellular patch clamp solution contained [in mM]: K-gluconate [140], EGTA [11], HEPES [10], and CaCl_2_ [1.32] and pH was adjusted to 7.3 using KOH. Target osmolarity was less than 290 mOsm.

Support solution was added consisting of ATP [4] and GTP [1]. More detailed methodology for this section can be found in ^31^.

### Cellular properties analysis

Cellular passive properties, action potential (AP) properties and firing properties were assessed in current clamp mode when cells were held at-70mV, and at nearly the same point in time after the electrode broke into the membrane. All values of absolute voltage, including resting membrane potential (RMP), AP absolute threshold (T) and overshoot voltage, and persistent inward currents (PICs) T, were adjusted by −10mV to approximately correct for the liquid junction potential. A more detailed analysis is available in our previous paper^19^. For the present study, all cellular properties were analyzed in Clampfit (Molecular Devices, RRID:SCR_011323).

#### Passive properties

In brief, a 3-second small hyperpolarizing current was injected into the tSPNs without triggering voltage dependent conductance to estimate cellular passive properties. Neuron input resistance (R_in_) was calculated by dividing maximal voltage deflection by the injected current. Although in non-isopotential neurons, the fit to voltage decay is a sum of exponential terms ^32^, the voltage decay to injected current in tSPNs exerts a single-exponential term, defined as the membrane time constant (τ_m_). Membrane capacitance (C_m_) was then calculated as τ_m_ divided by R_in_.

#### Action potential (AP) threshold and waveform

Three-second increasing depolarized current steps were applied to evaluate the AP properties and firing properties of tSPNs. The AP’s absolute threshold (T) was determined as the membrane potential at which the derivative of voltage, dV/dt, initiates an increase for the first AP. The relative T was measured as the potential difference between the absolute T and the membrane potential before the depolarizing step. Rheobase (I_rheo_) was estimated as the relative T divided by R_in_.

The AP waveform was assessed for the first AP at threshold. The overshoot voltage was measured as the peak voltage of the AP. Amplitude was defined as the voltage difference between the peak voltage and absolute T. The AP half-width represented the width of the spike at half of the AP amplitude.

#### Firing properties

All recorded tSPNs exhibited the capability of repetitive firing when subjected to depolarizing current steps. The maximum firing frequency was defined as the instantaneous frequency between the initial two spikes at each current onset, while the sustained firing frequency represented the mean instantaneous frequency for the last four spikes at each injected current. Frequency-current (ƒ-I) curve slope is the slope of the linear regression of the ƒ-I curve. The spike adaptation ratio was calculated as the ratio between the maximum and sustained firing rate at the maximum depolarizing current.

#### Sodium and potassium current properties

In voltage clamp mode, tSPNs were held at-90mV and 500 ms voltage steps were applied to all recorded tSPNs. Current traces were leak subtracted before analysis. Na current (I_Na_), A-type potassium current (I_A_), and potassium delayed outward rectifier (I_K_) were measured at-10mV. I_Na_ amplitude was measured as the maximum inward current within 2 ms of the voltage step onset and I_A_ as the peak outward current occurring 5-10 ms after the voltage step onset. I_K_ was the steady outward current at 490 ms of the voltage step.

#### Persistent inwards currents (PICs)

PICs were examined using triangular voltage ramp (-100mV to-10mV) with a 0.8 sec – 6 sec ramping up phase^33,34^ to eliminate sodium spikes. The PICs T was the absolute voltage when the PICs started to emerge. The PIC amplitude was measured as the peak inward current after leak subtraction.

#### Spontaneous excitatory postsynaptic current (sEPSC) properties

sEPSCs were recorded while tSPNs were held at –90mV for 12 seconds in voltage clamp. sEPSCs were visually identified, and those with amplitude below 1.5 times the noise amplitude were excluded from statistical analysis. Much larger currents indicative of spike dependent synaptic transmission were seen in 3/54 neurons, which were excluded from calculation of sEPSC frequency and amplitude. Tetrodotoxin (TTX, 1-2μM) was applied in 5 tSPN recordings to determine whether sEPSCs are action potential independent. sEPSC frequency was calculated as the total number of sEPSCs during this period divided by 12 seconds. Individual sEPSC amplitude was measured as the difference between the baseline and the sEPSC peak current. For each sEPSC, the decay phase from the peak was fitted with a single exponential function to calculate its time constant (τ_decay_). If two or more sEPSCs overlapped, only the amplitude of the first sEPSC was included in the statistical analysis, while the amplitude of subsequent sEPSCs and τ_decay_ values were excluded. Finally, individual amplitudes and τ_decay_ values were averaged for each tSPN to allow comparison among experimental groups, minimizing bias toward tSPNs receiving more frequent sEPSCs.

### TH^+^ tSPNs reconstruction

Experiments were conducted using sparse labelling TH^TdTomato^ mice.

#### Immunohistology for whole-mount thoracic sympathetic ganglia

Animals underwent intracardial perfusion with heparinized saline (0.9% NaCl, 0.1% NaNO_2_, 10 units/ml heparin), followed by 4% paraformaldehyde (0.5 M phosphate, 4% paraformaldehyde, NaOH for pH adjustment to 7.4). Immunohistology was processed at room temperature unless specified. The entire animal was post-fixed in 4% paraformaldehyde for 2 hours and subsequently transferred to a 30% sucrose solution, stored at 4 °C. Sympathetic ganglia were isolated and fasciae around the ganglia were carefully removed. The tissue was washed in 0.1 M PBS containing 0.3% Triton X-100 (PBS-T) overnight, followed by incubation in PBS-T solution 1 (80% PBS-T, 20% DMSO) overnight at 37°C, PBS-T solution 2 (79.7% PBS-T, 20% DMSO, 0.1% Tween20, 0.1% sodium deoxycholate, 0.1% Tergitol) overnight at 37°C, and PBS-T solution 3 (80% PBS-T, 20% DMSO, 0.3 M glycine) for 3 hours at 37°C. After DMSO treatment, tissues were washed in PBS-T (overnight followed by 4 x 1 hour) and then incubated for 5-7 days with primary antibodies. Subsequently, tissues were washed in PBS-T (6 x 1 hour), left in PBS-T overnight, and incubated with secondary antibodies for an additional 5-7 days.

After secondary antibodies incubation, tissues underwent washes in PBS-T (4 x 1 hour), followed by washes in 50 mM Tris-HCl (2 x 1 hour), and then left in 50mM Tris-HCl overnight. Finally, tissues were washed in 50mM Tris-HCl (2 x 1 hour) and allowed to dry before coverslipped.

#### Percentage of sparse labeled TH^+^ tSPNs in TH^TdTomato^ mice

Whole mount T5 ganglion (n = 4 naive mice, male, P > 34) were initially reacted for chicken anti-TH (1:250, Abcam, ab76442) and goat anti-Td (1:500, LSBio, LS-C340696) primary antibodies. Subsequently, the ganglion was reacted for Alexa 488 anti-chicken (1:100, Jackson Immunoresearch) and Cy3 anti-goat (1:250, Jackson Immunoresearch) secondary antibodies.

The imaging process utilized confocal microscopy (Olympus FV1000, 60x objective, oil immersion, 1.42NA, Z-step, <0.47 microns). For quantification, the number of TH^+^ tSPNs in T5 ganglion was determined using stereology with Stereoinvestigator (MBF Bioscience).

Concurrently, the number of Td^+^ neurons was manually counted using Fiji (ImageJ) software.

#### Td^+^ tSPNs reconstruction

Three weeks after SCI, animals (n = 3 mice, male, >P60) were perfused for experiments. Age-matched naïve mice (n = 3 mice, 2 males, 1 female) served as controls. Whole mount T4-T7 ganglia were initially reacted for goat anti-Td primary antibodies (1:500, LSBio, LS-C340696), followed by reactions for Cy3 anti-goat secondary antibodies (1:250, Jackson Immunoresearch). Images were captured using confocal microscopy (Olympus FV1000, 60x objective, oil immersion 1.42 or 1.35 NA, Z-step, <0.52 microns).

The cell bodies, axons, and dendrites of Td^+^ neurons (n = 22 neurons from SCI, n = 21 neurons from naive) were meticulously reconstructed using Neurolucida and NL360 software (MBF Bioscience). Only the neurons with neurites traceable to their termination were included in the statistical analysis. Quantitative measurement, performed with Neuroexplorer (MBF Bioscience), involved soma diameter and maximum cross section area (CSA), and number and length of primary/secondary axons and dendrites. In these measurements, an axon was defined as a neurite that exited a ganglion, traveled in an axon tract or was the single short projection from a cell without branching or significant change in diameter. A dendrite was defined as a neurite that was not an axon and extended for at least 5 microns from the cell surface. The primary neurite was the segment between the soma and the first branch point, while the secondary neurite encompassed the remaining dendritic extensions.

### *In-vivo* blood pressure measurement

Blood pressure was monitored using a high-throughput, non-invasive system (CODA®, Kent Scientific). Awake mice were individually secured in restraining tubes, allowing free tail movement. Each tube was placed on a heating pad to maintain body temperature above 36 C°. The mice remained calm throughout the recording sessions, with environmental interference minimized. An occlusion tail cuff was placed at the base of the tail to induce compression, while the remaining tail passed through a volume pressure recording sensor to detect systolic and diastolic pressure. Blood pressure was monitored weekly, starting two weeks before SCI or sham surgery and continuing for 3-6 weeks post-surgery. The two weeks of pre-surgery recordings were averaged and used as the baseline. Before each recording session, mice underwent a 20-minute daily acclimation period for two days, placed in the restraining tube on the heating pad without the tail cuff. On the recording day, a 15-minute acclimation period with the tail cuff on preceded the measurement. Blood pressure was recorded 15 times per session at one-minute intervals. The first five recordings were considered acclimation trials to familiarize the mice with tail cuff occlusion, and the subsequent ten recordings were averaged to determine blood pressure for the session. To minimize potential circadian effects, all acclimation and recording sessions were conducted at the same time each day. For experiments assessing blood pressure with and without bladder distension, initial recordings were taken with a full bladder. Immediately afterwards, the bladder was manually expressed, followed by an acclimation period and a second recording to measure blood pressure without bladder distension.

## Statistical analysis

This study employed a descriptive design, with statistical analyses conducted using SigmaPlot (Systat Software, RRID:SCR_003210). For tSPNs morphological, intrinsic and synaptic properties, significant differences were determined using either an unpaired Student’s t-test or the Mann-Whitney Rank Sum test when normality test failed, with P < 0.05 considered statistically significant. A *Z*-test was used to compare incidence rates between groups, also using a significance threshold of P < 0.05. For correlation analysis, Pearson’s Product-Moment Correlation Coefficient tests were performed to assess the correlation coefficient (*r*) and corresponding P-values. To control for multiple comparisons while maintaining an experiment-wise α of 0.05, a Šidák corrected α value (α_sid_) was calculated as:

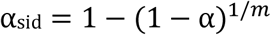

where m represents the number of comparisons. Correlations with P-values < α_sid_ and |*r*| > 0.7 were classified as strong correlations, while those with 0.4 < |*r*| < 0.7 were considered moderate. The coefficient of determination (*r*^2^) was calculated to assess the proportion of variance explained by the independent variable. In cases where parameter pairs exhibited moderate correlation (|*r*| > 0.4) but failed to reach significance as P-values < α_sid_ due to intrinsic variability, they were still reported as moderate correlations, with the caution that their interpretation should be approached carefully.

### Computational model

We updated a previously described conductance-based computational model of mouse tSPNs^19^ to effectively describe experimental recordings for an individual neuron simultaneously tuned across different electrophysiological modalities, including voltage-clamp (VC) step and ramp, as well as current clamp (CC) protocols. Relevant ion channels were identified based on previously studied mRNA profiles in tSPNs^28^ and the maximal conductances, decay time constants, and other kinetic properties of these ion channels were determined based on experimental data. In particular, we modeled the dynamics of fast TTX sensitive Na^+^, high-threshold long lasting Ca^2+^, and A-type K^+^ currents based on the equations derived from rat nodose sensory neurons^35^, with the model parameters for the transient Na^+^ and A-type K^+^ currents modified using VC step protocol data, further separated by the application of the Na^+^ channel blocker TTX. Various long-lasting currents (delayed rectifier K^+^, M-type K^+^, Ca^2+^ dependent K^+^ with current equations unmodified from previous model^19^) were separated based on their characteristic dynamics seen under VC ramp protocol, and their aggregate under VC step protocol. Using data from CC recordings, all channel properties were also simultaneously tuned to match the passive and firing properties of the cells in response to current injections. A model of the electrode resistance and capacitance^36^ was included to separate experimental artifacts from ion channel dynamics and to describe the voltage dependent delays observed in the onset of inward Na^+^ currents under VC step protocol. We included a Na^+^/K^+^ pump current with cubic dependency on [Na^+^] ion concentration^37^, updated the Nernst potentials for Na^+^, K^+^, Ca^2+^ channels based on membrane ion concentrations, added an h-current channel, a leak channel conducting a combination of Na^+^ and K^+^ currents, and updated their temperature dependence (temperature kept constant at 296K for this study). Detailed equations describing the model as well as parameter detail differences between representative sham and SCI cells are provided in the **Supplementary Methods** section.

Two models were used in this study. The model relating to tSPN recruitment (**Fig 3C**, **Fig 8D**) incorporated the average passive membrane properties measured experimentally in the sham or SCI groups, reflecting their physiological integrative capability. For the *f*-I curves simulations (**Fig 7A_4_-7B_4_**), parameters were first optimized to replicate the average firing properties of tSPNs in the sham group, serving as the sham model neuron. These parameters were then adjusted based on the experimentally observed ratio changes between sham to SCI group to generate the SCI model neuron.

**Figure 3.**
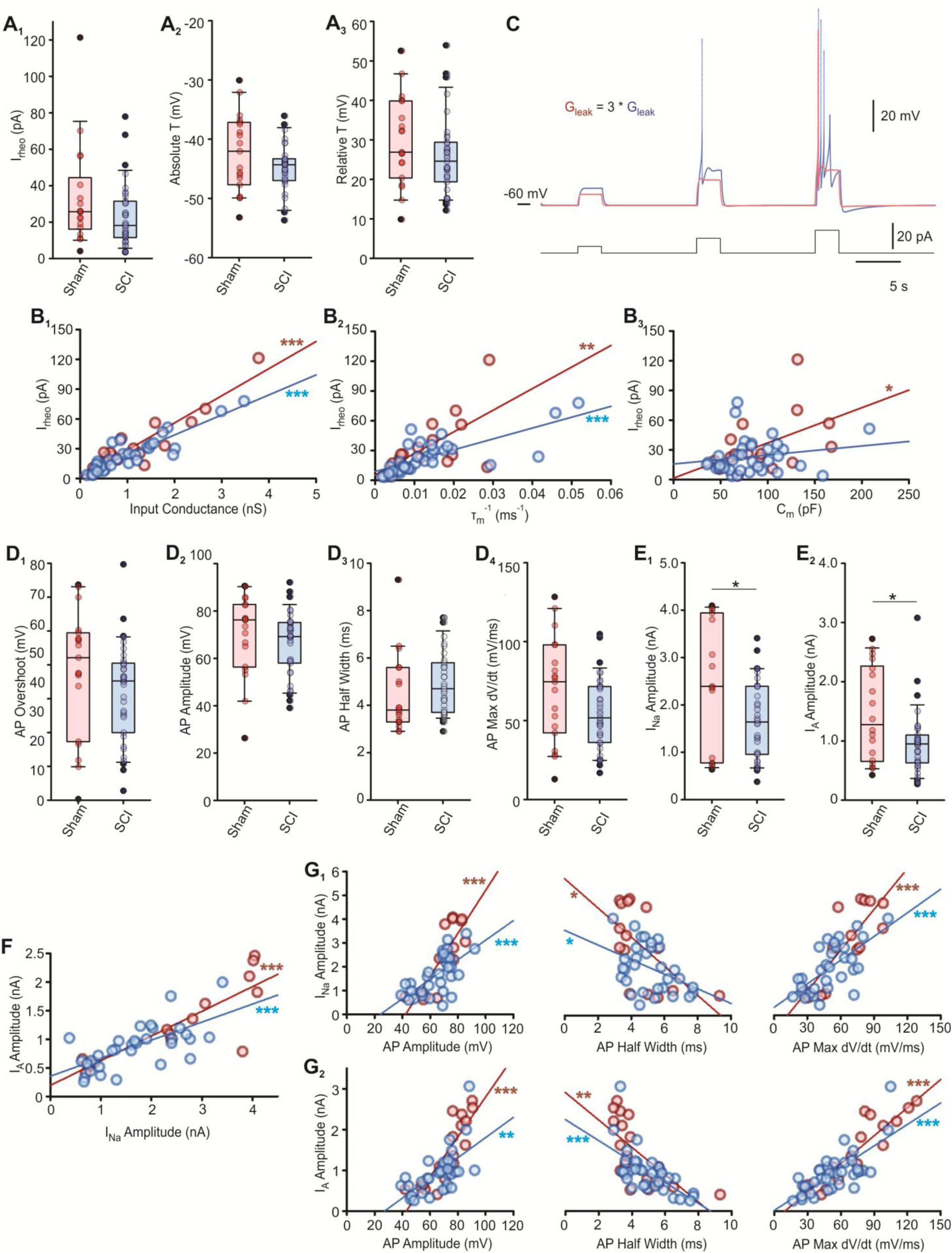
Whole population tSPNs action potential and current properties. Comparison of whole population tSPNs threshold, spike wave form and current properties in sham and SCI groups. ***A,*** Summary of whole population tSPNs threshold properties. No significant difference between sham and SCI group in rheobase (***A_1_,*** I_rheo_, P = 0.13), absolute threshold (***A_2_,*** Absolute T, P = 0.28), and relative threshold (***A_3_,*** Relative T, P = 0.20), unpaired student t-test. ***B,*** Correlations between I_rheo_ and passive properties. ***B_1_,*** Strong correlation between I_rheo_ and input conductance (R_in_^-1^) in both sham (r = 0.92, P = 5.8×10^-8^) and SCI group (r = 0.90, P = 9.3×10^-14^). ***B_2_,*** Moderate correlation between I_rheo_ and inverse of time constant (τ_m_^-1^) in sham group (r = 0.63, P = 6.6×10^-3^), but strong correlation in SCI group (r = 0.74, P = 4.1×10^-7^). ***B_3_,*** Moderate correlation between I_rheo_ and C_m_ in sham group (r = 0.49, P = 0.046), but not in SCI group (r = 0.18, P = 0.29). Pearson’s Product-Moment Correlation test. ***C,*** Voltage response of a model cell when simulating the increase in R_in_ after SCI. The leak conductance in the red trace is 3-fold greater than that in the blue trace, analogous to the approximate mean R_in_ values in sham and chronic tSPNs, respectively (1436MΩ in red; 2100MΩ in blue). A series of 3-second incrementing injected current steps were given to the model cell. Less current was required to trigger action potential, representing a reduction in I_rheo_. ***D,*** Summary of whole population tSPN spike waveform. No significant difference in action potential overshoot (***D_1_,*** P = 0.21), amplitude (***D_2_,*** P = 0.20), half width (***D_3_,*** P = 0.07) and rising slope (***D_4_,*** max dV/dt, P = 0.05). ***E,*** Summary of whole population tSPN sodium and A-type potassium current properties. Significant reduction in sodium current (***E_1_,*** I_Na_, P = 0.023) and A current (***E_2_,*** I_A_, P = 0.047) amplitude after SCI, unpaired student t-test. ***F,*** Strong correlation between I_Na_ and I_A_ amplitude in sham group (r = 0.86, P = 3.85×10^-5^), and moderate correlation in SCI group (r = 0.67, P = 3.12×10^-5^). ***G,*** Correlations between I_Na_ and I_A_ amplitude and spike waveform. ***G_1_,*** Strong positive correlation between action potential amplitude and rising slope with I_Na_ amplitude in sham group (r = 0.83, P = 1.3×10^-4^, and r = 0.84, P = 8.9×10^-5^ respectively), but moderate positive correlation with I_Na_ amplitude in SCI group (r = 0.65, P = 7.7×10^-5^, and r = 0.67, P = 4.2×10^-5^ respectively). Moderate negative correlation between action potential half width and I_Na_ amplitude in both sham (r =-0.61, P = 0.016) and SCI group (r =-0.41, P = 0.024). ***G_2_,*** Strong positive correlation between action potential amplitude with I_A_ amplitude in sham (r = 0.85, P = 9.8×10^-6^), but moderate in SCI group (r = 0.62, P = 1.1×10^-4^). Moderate negative correlation between action potential half width and I_A_ amplitude in both sham (r =-0.68, P = 1.8×10^-3^) and SCI group. (r =-0.63, P = 5.8×10^-5^). Strong positive correlation between action potential rising slope and I_A_ amplitude in both sham (r = 0.90, P = 3.9×10^-7^) and SCI group. (r = 0.73, P = 1.1×10^-6^). Pearson’s Product-Moment Correlation test. *P < 0.05, ** P < 0.01, *** P < 0.001

**Figure 4.**
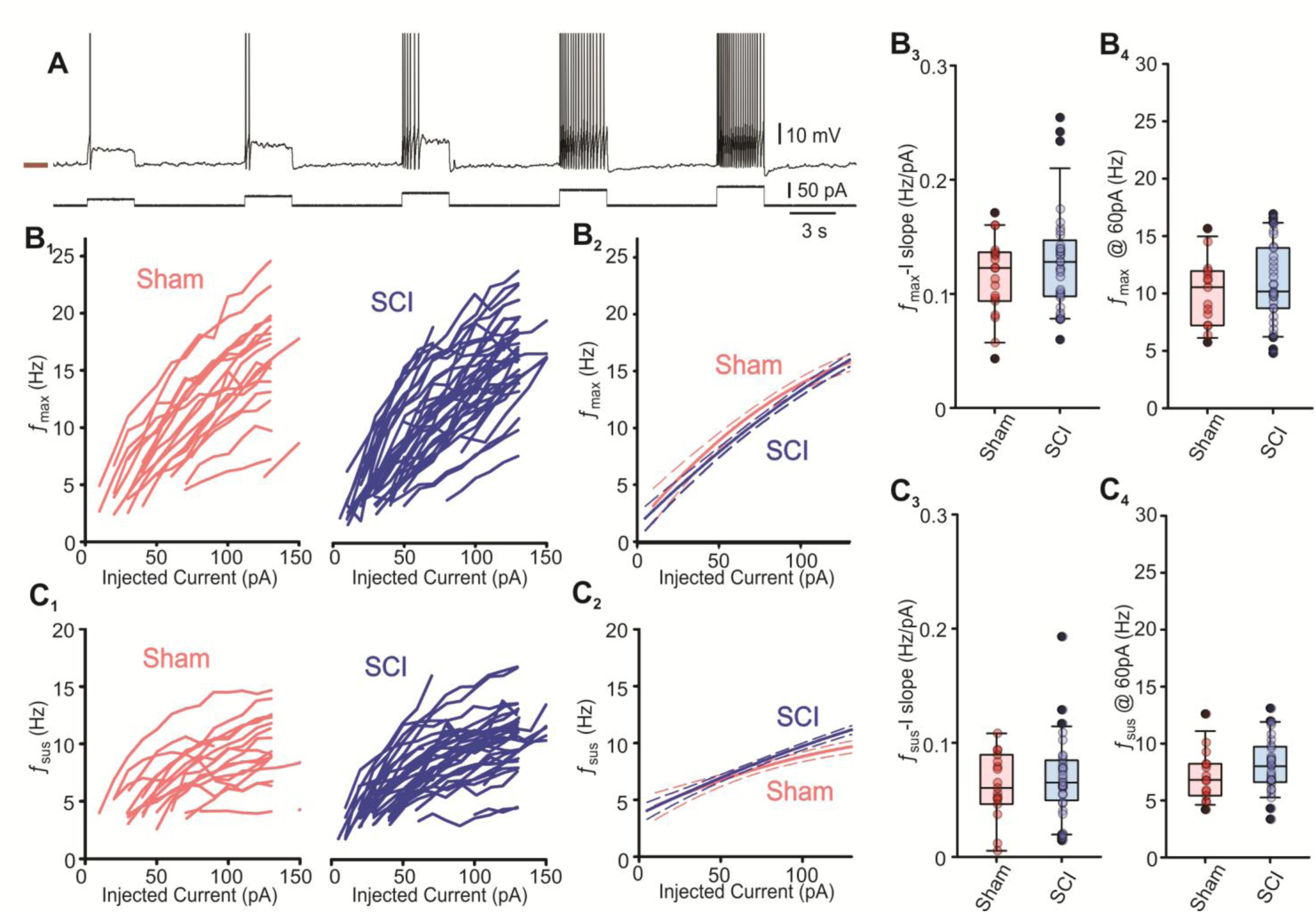
Whole population tSPN firing properties. Comparison of whole population tSPNs repetitive firing properties in sham and SCI groups. ***A,*** Example trace from a current clamp recording showing increased firing frequency in response to incrementing current steps. Action potential is truncated. ***B-C,*** Maximum instantaneous firing properties and sustained firing properties. ***B_1_-C_1_,*** Maximum instantaneous firing frequency (*f*_max_) and sustained firing frequency (*f*_sus_) is plotted versus injected current (*f*_max_-I curve and *f*_sus_-I curve, respectively) for each individual tSPN. ***B_2_-C_2_,*** Polynomial best fit (solid line) of sham (red) and SCI (blue) *f*_max_-I curves in B_1_ and *f*_sus_-I curves in C_1_. Dash lines represent 95% confidence intervals. ***B_3_-C_3_,*** No significant difference between sham and SCI in *f*_max_-I curve slope (P = 0.41) and *f*_sus_-I curve slope (P = 0.63**). *B_4_-C_4_,*** No significant difference between sham and SCI in *f*_max_ (P = 0.422) and *f*_sus_ (P = 0.247) when injected with a 60pA current, unpaired student t-test.

**Figure 5.**
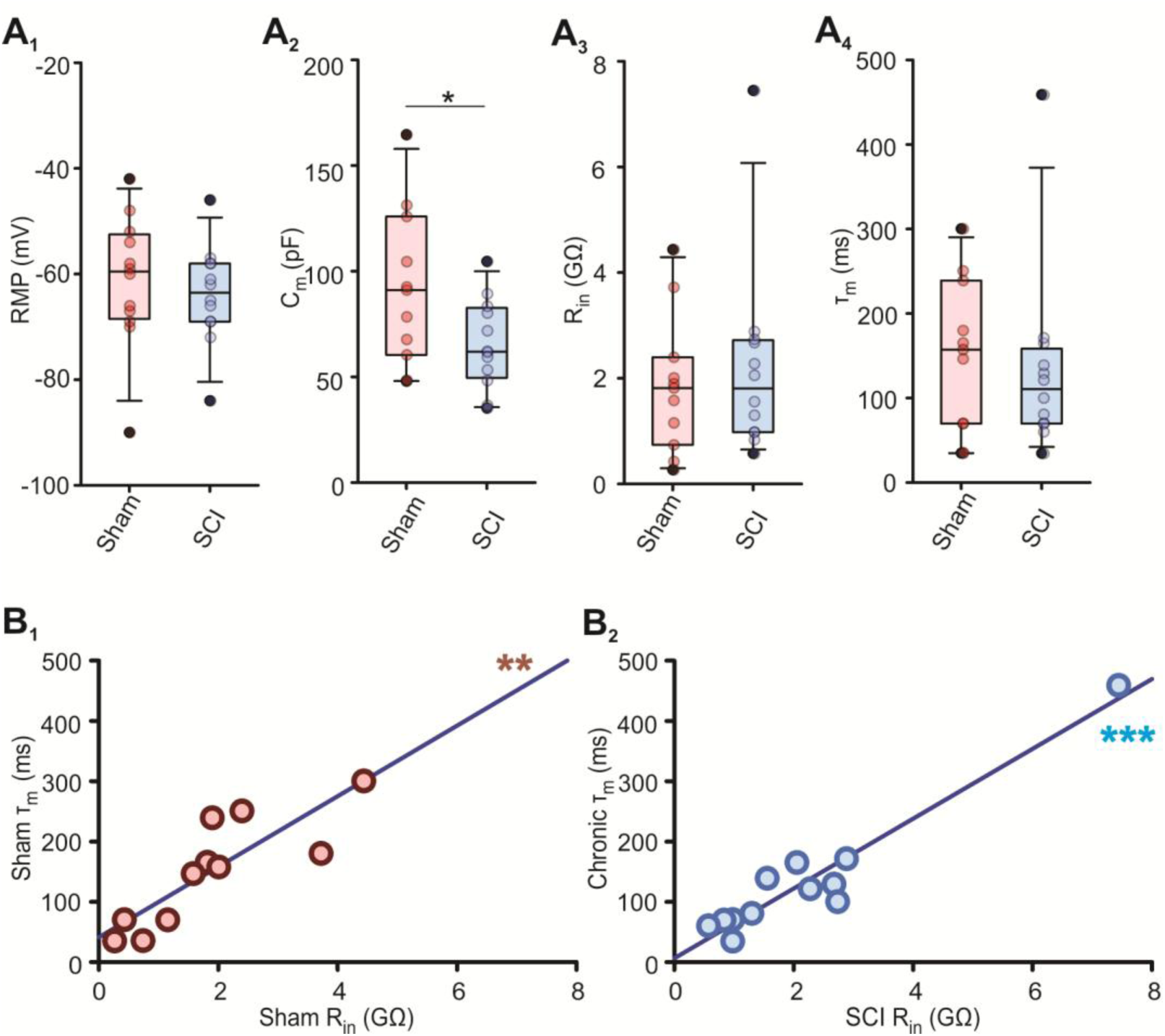
NPr^+^ tSPN passive properties. Comparison of NPY^+^ tSPNs passive properties in sham and SCI groups. ***A,*** Summary of passive properties of NPY^+^ tSPNs in sham (n = 6 mice, pink) and SCI (n = 8 mice, blue) groups. Membrane capacitance was significantly decreased after SCI (***A_2_,*** C_m_, P = 0.045). No significant difference was observed in resting membrane potential (***A_1_,*** RMP, P = 0.56), input resistance (***A_3_,*** R_in_, P = 0.67) and time constant (***A_4_***, τ_m_, P = 0.39), unpaired student t-test. Box plots and horizontal bar within represent the interquartile range and median, respectively. Error bars extend to the most extreme data point that is within 1.5 times the interquartile range. ***B,*** R_in_ was strongly correlated with τ_m_ in both sham (***B_1_,*** r = 0.84, P = 1.11×10^-3^) and SCI groups (***B_2_,*** r = 0.96, P = 8.37×10^-7^). Solid line indicates linear simple regression, Pearson’s Product-Moment Correlation test. *P < 0.05, ** P < 0.01, *** P < 0.001.

**Figure 6.**
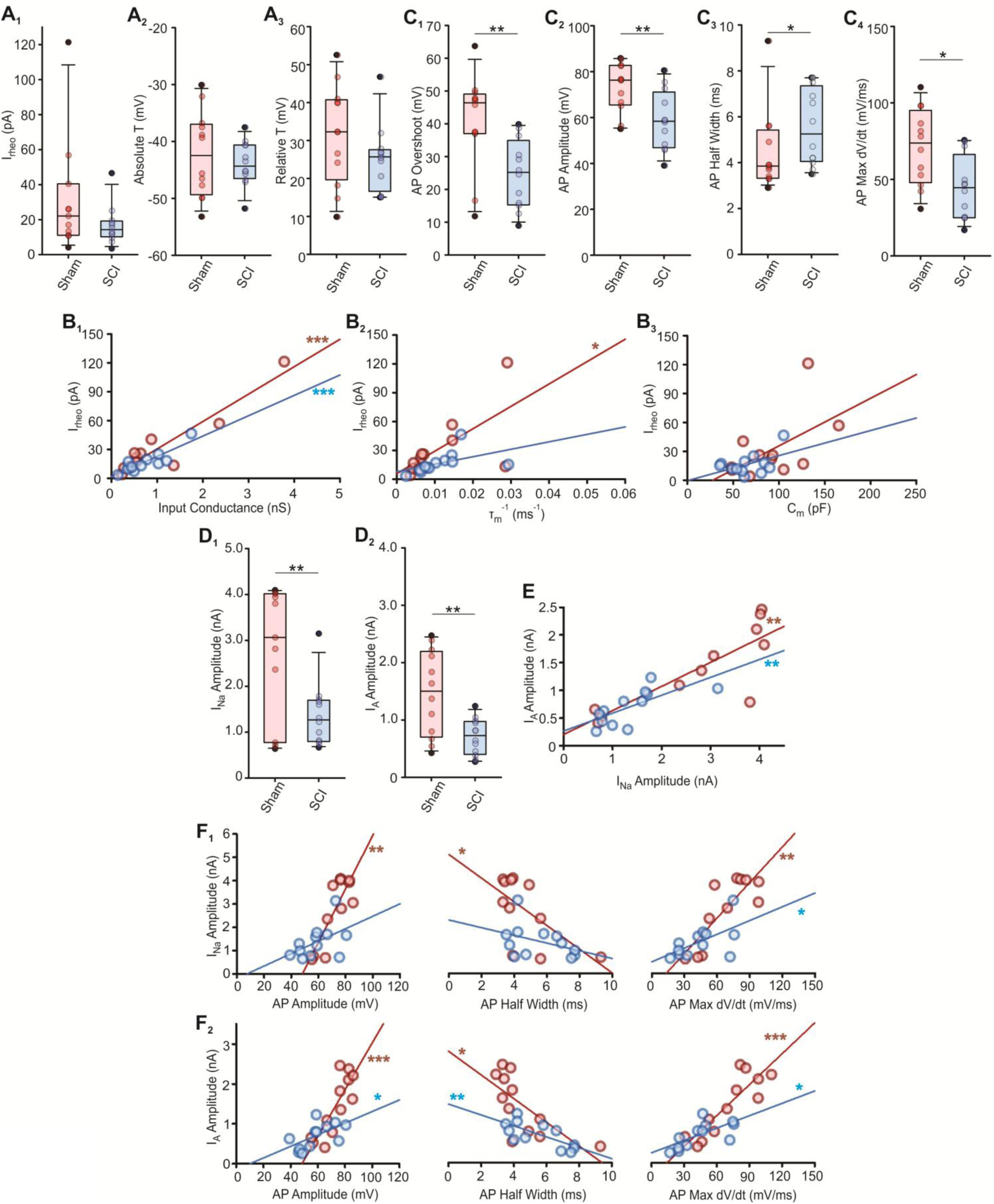
NPY^+^ tSPN threshold, spike waveform and current properties. Comparison of NPY^+^ tSPNs action potential threshold, waveform and current properties in sham and SCI groups. ***A,*** Summary of spike threshold properties. No significant difference between sham and SCI group in rheobase (***A_1_,*** I_rheo_, P = 0.20), absolute threshold (***A_2_,*** P = 0.52) and relative threshold (***A_3_,*** P = 0.18), unpaired student t-test. ***B,*** Correlations between I_rheo_ to passive properties. ***B_1_,*** Strong correlation between I_rheo_ and input conductance in both sham (r = 0.93, P = 2.51×10^-5^) and SCI group (r = 0.88, P = 1.61×10^-4^). ***B_2_,*** Moderate correlation between I_rheo_ and inverse of time constant (τ_m_^-1^) in sham group (r = 0.66, P = 0.028) and SCI group (r = 0.49, P = 0.10). ***B_3_,*** No correlation between I_rheo_ and C_m_ in either sham (r = 0.55, P = 0.08) or SCI group (r = 0.50, P = 0.10, respectively). Pearson’s Product-Moment Correlation test. ***C,*** Summary of NPY^+^ tSPN action potential waveform. Significant reduction in action potential overshoot potential (***C_1_,*** P = 0.005), amplitude (***C_2_,*** P = 0.006) and maximum rising slope (***C_4_,*** max dv/dt, P = 0.01). ***C_3_,*** Significant increase in action potential half width (P = 0.046). Unpaired student t-test. ***D,*** Significant reduction in I_Na_ current amplitude (***D_1_,*** P = 0.007) and I_A_ (***D_2_,*** P = 0.004) after SCI, unpaired student t-test. ***E,*** Strong correlation between I_Na_ and I_A_ amplitude in sham group (r = 0.83, P = 1.46×10^-3^), but moderate correlation in SCI group (r = 0.71, P = 9.83×10^-3^). ***F,*** Correlations between I_Na_ and I_A_ amplitude and spike waveform. Strong correlation between action potential amplitude with I_Na_ and I_A_ amplitude in sham group (r = 0.83, P = 1.45×10^-3^, and r = 0.83, P = 8.07×10^-4^ respectively), but moderate to no correlation in SCI group (r = 0.50, P = 0.099, and r = 0.59, P = 0.042 respectively). Moderate correlation between action potential half width with I_Na_ and I_A_ amplitude in sham group (r =-0.63, P = 0.038, and r = - 0.70, P = 0.012 respectively), but moderate to no correlations in SCI group (r =-0.40, P = 0.203, and r =-0.72, P = 8.31×10^-3^ respectively). Moderate correlation between action potential rising slope with I_NA_ amplitude in both sham group (r = 0.82, P = 2.05×10^-3^) and SCI group (r = 0.59, P = 0.044). Strong correlation between action potential rising slope and I_A_ amplitude in sham group (r = 0.87, P = 3.63×10^-4^), but moderate in SCI group. (r = 0.68, P =0.015). Pearson’s Product-Moment Correlation test. *P < 0.05, **P < 0.01, *** P < 0.001

**Figure 7.**
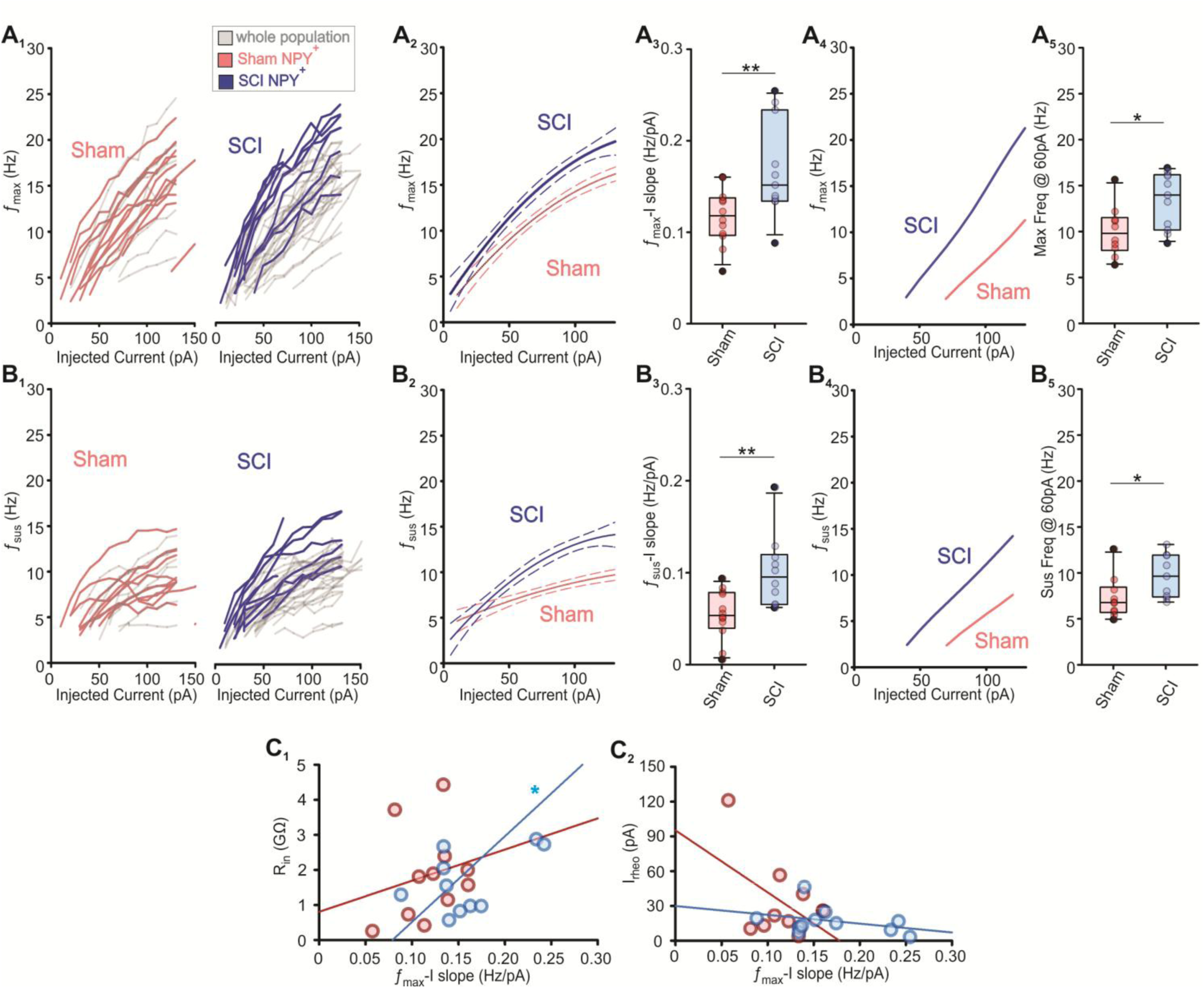
NPY^+^ tSPNs repetitive firing properties. Comparison of repetitive firing properties in NPY^+^ tSPNs in sham and SCI groups. ***A-B,*** Maximum instantaneous firing properties (*f*_max_) and sustained firing properties (*f*_sus_). ***A_1_-B_1_,*** *f*_max_ and *f*_sus_ are plotted versus injected current (*f*_max_-I curve and *f*_sus_-I curve respectively) for each individual NPY^+^ tSPN. Colored lines indicate NPY^+^ tSPNs. Gray lines indicate the whole population tSPNs. ***A_2_-B_2_,*** Polynomial best fit (solid line) of sham (pink) and SCI (blue) *f*_max_-I curves in A_1_ and *f*_sus_-I curves in B_1_. Dash line represents 95% confidence intervals. ***A_3_-B_3_,*** Significant increase in *f*_max_-I curve slope (P = 0.009) and *f*_sus_-I curve slope (P = 0.004) in SCI group compared to sham. ***A_4_-B_4_,*** Simulated *f*_max_-I and *f*_sus_-I curve by simulating mean value ratio changes in NPY^+^ tSPNs C_m_, R_in_, I_NA_, I_A_ and I_K_ from sham and SCI groups. ***A_5_-B_5_,*** Significant increase after SCI in *f*_max_ (P = 0.015) and *f*_sus_ (P = 0.039) when NPY^+^ tSPNs injected with 60pA current. Unpaired student t-test. ***C,*** Correlations between *f*_max_-I curve slope with R_in_ and I_rheo_. ***C_1_,*** Moderate correlation between *f*_max_-I curve slope and R_in_ was observed in SCI group (r = 0.67, P = 0.024), but not in sham (r = 0.22, P = 0.52). ***C_2_,*** *f*_max_-I curve slope was not correlated with I_rheo_ in sham group (r =-0.51, P = 0.11), or after SCI (r =-0.35, P = 0.29). Pearson’s Product-Moment Correlation test. *P < 0.05, ** P < 0.01.

**Figure 8.**
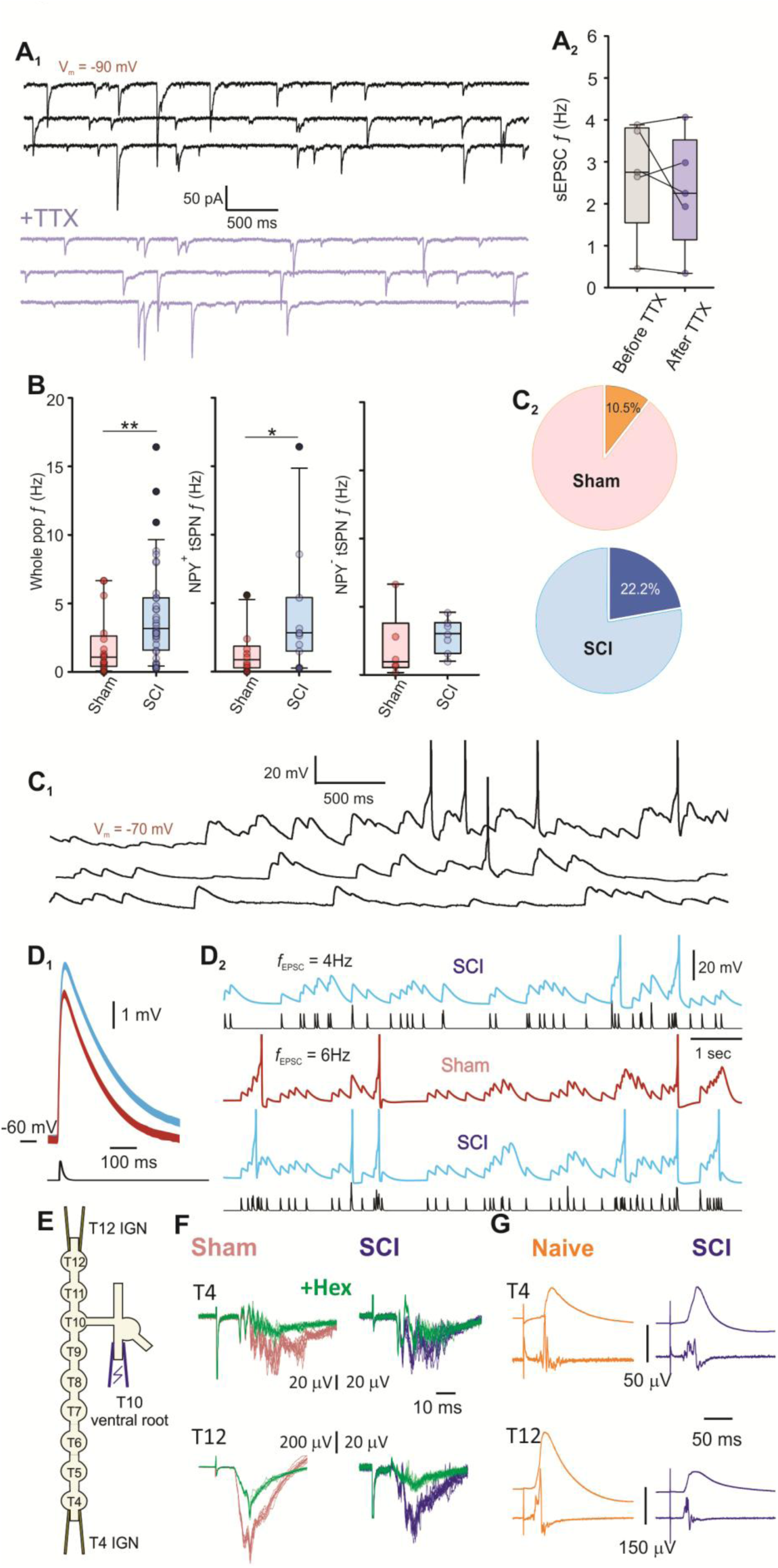
Whole population and NPY^+^ tSPN sEPSC properties. Comparison of sympathetic preganglionic spontaneous inputs to tSPNs in sham and SCI groups. ***A,*** sEPSCs are action potential-independent quantal events. **A_1,_** Examples of sEPSCs from a tSPN in a SCI mouse. sEPSCs were resistant to action potential dependent transmission block with TTX (2 μM). V_m_ was held at-90mV. ***A_2_***, sEPSC frequency was unchanged with TTX (1-2 μM, n = 5, P = 0.68). ***B,*** Summary of sEPSC frequency in the whole population, NPY^+^ and NPY^-^ tSPNs. Significant increase in sEPSC frequency after SCI compared to sham in whole population (P = 0.007) and NPY^+^ tSPNs (P = 0.038). A trend of increase in sEPSC frequency in NPY^-^ tSPNs (P = 0.472). ***C,*** sEPSPs are capable of summating to initiate action potentials. ***C_1_***, Spontaneous EPSPs display capacity for temporal summation sufficient for spike recruitment. (V_m_ was held at-70mV and spikes are truncated to emphasize synaptic events). ***C_2_***, Incidence of tSPNs capable of spontaneous spiking when membrane potential was held at-70mV in sham (2/19) and SCI groups (8/36) was not statistically different (P = 0.28; z-test). ***D,*** Model tSPNs spiking in response to spontaneous synaptic inputs. The model neurons adopted the average passive membrane properties measured experimentally from sham and SCI NPY^+^ tSPNs. 40pA randomly timed synaptic currents (black traces in D_2_) were given to both model neurons. ***D_1_***, Unitary EPSP in SCI model neuron was greater than the EPSP in sham model neuron. ***D_2_***, Both sham and SCI model neurons (RMP =-60mV) generated spikes (truncated in Fig) in response to spontaneous synaptic currents, with EPSPs summation leading to more spiking for synaptic currents delivered at an average frequency of 6Hz compared to 4Hz. The SCI model neuron exhibited more frequent spiking compared to the sham model. ***E,*** Schematic of *ex vivo* thoracic chain preparation for population compound potential recording. Suprathreshold electrical recruitment (200 μs, 200 μA) of T10 preganglionic axons was achieved with suction electrodes on the ventral root. Evoked responses were recorded from suction electrodes placed on the cut end of the interganglionic nerves (IGN) at rostral T4 and caudal T12. ***F,*** Example traces of electrically evoked compound action potentials (CAPs) at T4 and T12 IGN. Shown are 10 raw events overlaid both before (orange traces in sham, blue traces in SCI) and after (green traces) bath applied hexamethonium (hex; 50 – 100 μM). The remaining CAPs after hex represent the preganglionic component of the evoked response. ***G,*** Electrical recruitment of preganglionic axons from the T10 ventral root evokes synaptic responses from suction electrodes attached at T4 and T12 interganglionic nerves adjacent to respective sympathetic ganglia. Volley is filtered at 50 Hz high-pass while population synaptic response is filtered at 15 Hz low-pass. Note the presence of post-EPSP hyperpolarization presumably due to activation of an M current reduces EPSP duration. Responses shown are averages of 10 events. Evoked responses at T12 are significantly smaller in the SCI population due at least in part to significant reduction in the preganglionic volley.

## Results

### T2 SCI leads to significant hemodynamic alterations

Blood pressure measurements were conducted weekly, beginning two weeks before sham or SCI surgery to establish baseline values and continuing for 3 to 6 weeks post-surgery. Post-surgery blood pressure values were normalized to each mouse’s baseline. In SCI mice, systolic, diastolic, and mean pressure significantly decreased during the first two weeks post-surgery, followed by a subsequent recovery to pre-surgery levels. In contrast, these parameters remained unchanged after sham surgery (Fig 1B, Sup Fig 1A-B). Bladder distension is a major trigger for AD, and an increasing blood pressure is typically accompanied by a slower heart rate during AD^38^. To further investigate the development of AD, blood pressure and heart rate were compared with or without bladder distension at 3-6 weeks post-SCI. Systolic pressure and diastolic pressure showed significant increases of 9.6% and 10.6%, respectively, while heart rate exhibited a numerical 9.4% decrease with bladder distention (Sup Fig 1C). These findings suggest hemodynamic alterations have developed at the chronic stage of SCI (≥ 3 weeks po-SCI) in our experimental model.

### When not differentiating sub-populations, tSPNs showed a trend of cell shrinkage and an increase in integrative capacity at the chronic stage of SCI

To investigate intrinsic plasticity in tSPNs at the chronic stage of SCI, we first assessed tSPN properties without differentiating between the 7 functional groups that have been identified by single-cell RNA sequencing studies ^28^. We refer to this undifferentiated dataset as whole population tSPNs. Most recordings were conducted in T3-T5 thoracic sympathetic ganglia (**Fig 1C**). **Table 1** summarizes the intrinsic properties of whole population tSPNs in both sham and chronic SCI groups. Note that passive and active membrane properties measured in sham mice were comparable to previously obtained data in naive mice^19^.

**Table 1.**
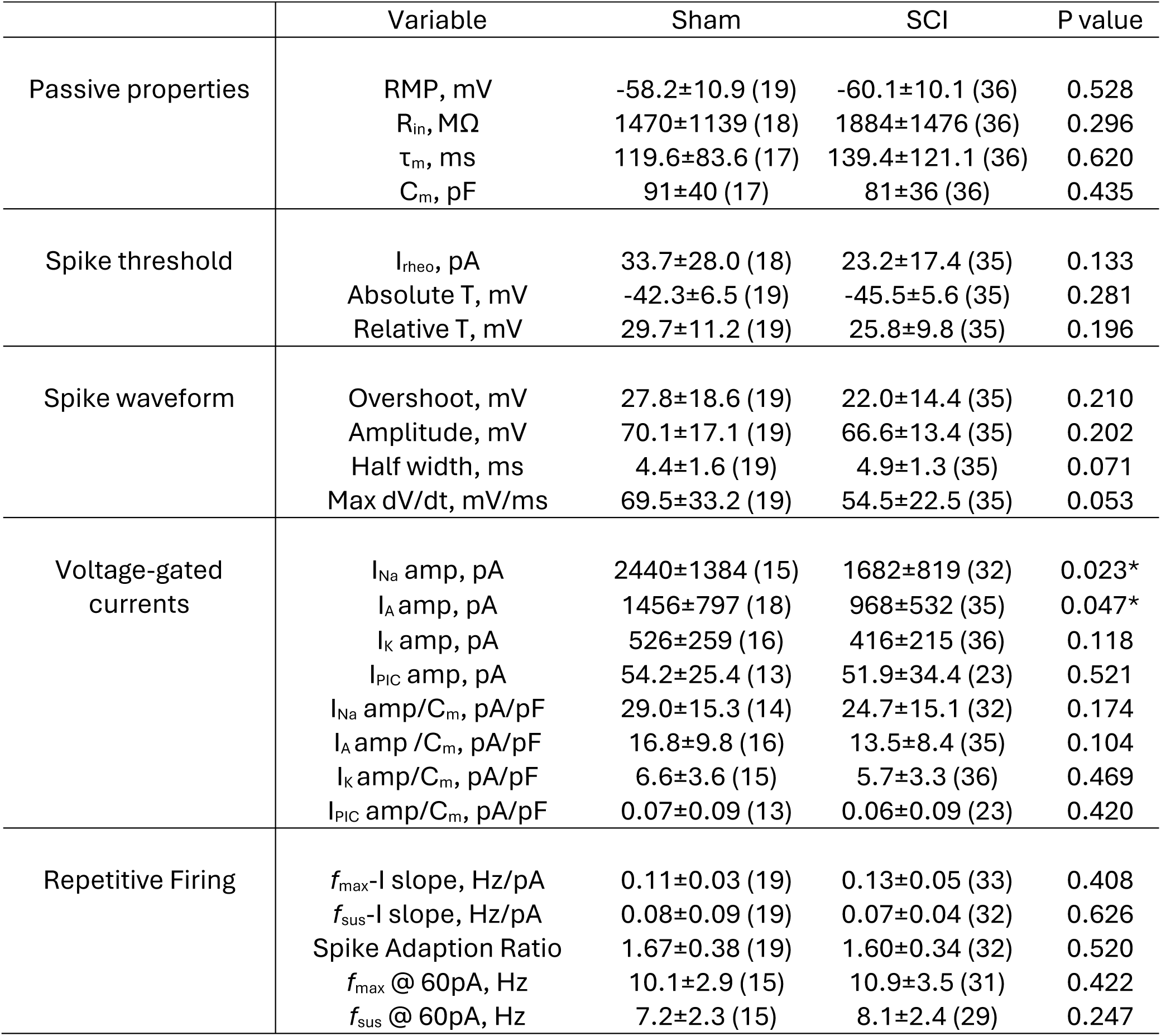
The whole population tSPNs intrinsic properties in Sham and SCI groups. Intrinsic properties of the whole population tSPNs. Values of basic properties of whole population tSPNs in sham and SCI groups. Data is represented as mean ± SD (number of observations). **RMP**, resting membrane potential; **R_in_**, input resistance; **τ_m_**, time constant; **C_m_**, capacitance; **I_rheo_**, rheobase; **T**, threshold; **Max dV/dt**, action potential maximum rising slope; **I_Na_**, sodium current; **I_A_**, A-type potassium current; **I_K_**, potassium delayed outward rectifier; **I_PIC_**, persistent inward current; **amp**, amplitude; ***f*_max_**, maximum instantaneous frequency; ***f*_sus_**, sustained firing frequency; **I**, current; *P<0.05. Unpaired student t-test.

Resting membrane potential (RMP), cell size (capacitance, C_m_), and passive membrane properties (input resistance R_in_ and time constant τ_m_) are important to synaptic integration. We noted a numerical 28.2% increase in neuron R_in_ (**Fig 1D_1_**) and a numerical 16.6% increase in τ_m_ (**Fig 1D_2_**). Both measured parameters exhibited a >10-fold range in both sham and SCI groups. Neuron C_m_ (**Fig 1D_3_**) values varied by a > 4-fold range and numerically decreased by 11.0% after SCI. RMP (**Fig 1D_4_**) showed minimal change after SCI. Overall, in both sham and SCI groups, a wide range of passive membrane properties was observed, and no significant differences were evident after SCI. Consistent with previous findings in naive tSPNs ^39^, we observed a strong correlation between R_in_ and τ_m_ in both sham and chronic SCI groups (**Fig 1E**). However, no significant correlation was found between R_in_ and C_m_, indicating that differences in tSPN R_in_ are governed by variation in resistivity rather than cell size. A summary of correlations among whole population tSPNs measured parameters is provided in **Sup Table 1**.

We utilized a sparse labeling TH^Cre^ line ^27^ crossed with TdTomato reporter mice to reconstruct the morphology of a presumed representative sample of adrenergic tSPNs (**Fig 2A**). In T5 ganglion from four animals, TH^TdTomato^ neurons constituted 1.6 - 5.9% (mean ± SD: 3.4 ± 1.9) of the TH^+^ tSPN population. Corresponding to the substantial variability in cell surface area estimated with C_m_, reconstructed tSPNs exhibited a wide range in soma size, number of neurites, and complexity of neurites within the same ganglion (**Fig 2B**). While soma diameter remained unchanged after SCI, primary dendrite length was numerically reduced by 30.7% (**Fig 2C_1_**), and secondary dendrite length significantly decreased by 37.1% (**Fig 2C_2_**) in mice 3 weeks post-SCI (**Sup Table 2**). Thus, even within this extensive variability in cell morphology, arborizations were significantly reduced. Detailed measurements for anatomical parameters of tSPNs are summarized in **Sup Table 2**.

### Overall tSPN recruitment is driven by variation in R_in_ in both sham and SCI populations

To explore alterations in tSPNs’ excitability after SCI, we determined the minimum depolarizing current required to initiate spiking (rheobase, I_rheo_) and measured its relative voltage threshold (magnitude of depolarization to trigger action potential, relative T) and absolute threshold (membrane voltage when action potential was initiated, absolute T). Assuming the relative threshold obeys Ohm’s law, the observed numerical increase in R_in_ after SCI should lead to a reduction in I_rheo_. Indeed, an overall 28.2% increase in R_in_ led to a numerical 31.2% decrease in I_rheo_ (**Fig 3A_1_**). Moreover, variation in R_in_ stood out as the single best-correlated indicator of excitability, surpassing τ_m_ and C_m_ (**Fig 3B**). Like R_in_, I_rheo_ values varied over a >10 fold range.

The decreased slope of this relation after SCI (**Fig 3B_1_**) suggests that other factors contribute to an excitability increase after SCI, which may include slight, albeit not significant, reductions in voltage thresholds (↓13.1% in relative T and ↓7.6% in absolute T) (**Fig 3A_2-3_**). The significant correlations of I_rheo_ to R_in_ and τ_m_ but not C_m_ strongly support membrane resistivity as a prominent mechanism guiding variability in excitability across tSPN population ^19,40^. To further verify this finding, we established a tSPN computational model with intrinsic property parameters that fall within the range of experimental measurements observed in tSPNs from the sham group. When we decreased the leak conductance to mimic the increase in R_in_ after SCI, I_rheo_ was reduced as expected (**Fig 3C**).

### Overall, sodium and A-type potassium current densities and spike waveforms were unchanged after SCI

Next, we examined the action potential properties of the whole population tSPNs after SCI. There was no significant difference in spike waveform measures (spike height, rising slope and half width) (**Fig 3D**). The conductances expected to govern spike shape in sympathetic neurons are the voltage-dependent sodium inward current (I_Na_) and the fast-activating, fast-inactivating A-type potassium outward current (I_A_) ^41^. The measured magnitudes of I_Na_ and I_A_ varied over a 4-fold range and were strongly correlated in both sham and SCI mice (**Fig 3E-F**). Both I_Na_ and I_A_ were significantly and comparably decreased in amplitude after SCI (31.1% and 33.5%, respectively; **Fig 3E**), presumably reflecting a tight molecular co-regulation on spike waveform. Variation in spike amplitude, rising slope and half-width were correlated with both I_Na_ and I_A_ amplitude in both sham and SCI groups (**Fig 3G**). Interestingly, current densities obtained by normalizing I_Na_ or I_A_ to C_m_ removed this significant reduction (**Table 1**), suggesting that channel densities are conserved in the presence of cell shrinkage after SCI to preserve spike shape.

### Overall, tSPNs repetitive firing properties are highly resilient to SCI

Repetitive firing properties were characterized via a series of injected current steps. All tSPNs fired repetitively and underwent spike frequency adaptation in response to injected current steps in both sham and chronic SCI groups (**Fig 4A**). The relationship between output frequency (*f*) and injected depolarizing current (I) is commonly assessed with *f*-I curves ^39,42^. Overlaid plots of individual *f*-I curves of maximal initial (*f*_max_) and sustained (*f*_sus_) firing frequency depict the large range in cellular excitability (**Fig 4B_1_, C_1_**). Polynomial best fits of sham and SCI *f*-I curves overlapped (**Fig 4B_2_, C_2_**). Graphically, SCI mice *f*-I curves had slight leftward shifts, higher slopes, and the highest values. While such differences would support an SCI-induced increase in tSPN excitability, these changes were not statistically significant (**Fig 4B_3-4_, C_3-4_**), indicative of an unchanged output gain after SCI.

Population maximum (*f*_max_-I) and sustained (*f*_sus_-I) *f*-I slopes were not strongly correlated with any individual passive or active membrane property nor underlying voltage-gated conductances (**Sup Table 1**). This suggests that *f*-I slope regulation is multifactorial ^19^. Different dependencies in composite subpopulations may also mask interactions.

### Studies in NPY^+^ tSPNs

tSPNs are functionally comprised of seven molecularly distinct populations ^28^. The wide range in excitability observed in whole population tSPNs may indicate the presence of various functional subgroups of tSPNs with more restricted ranges in membrane properties, as seen with cell soma ^28^. Moreover, SCI may induce distinct effects in individual populations, and these effects may be obscured by the absence of or opposite changes in other populations. Therefore, we selectively examined the effects of SCI on NPY^+^ tSPNs using NPY^TdTomato^ mice. Approximately 50% of tSPNs express NPY ^16,28^, the majority (80%) of which innervate blood vessels ^43^.

### (A) SCI induces cell shrinkage in NPY^+^ tSPNs

**Table 2** summarizes the intrinsic properties of NPY^+^ tSPNs in both sham and chronic SCI groups. Passive properties of NPY^+^ tSPNs were compared between sham and chronic SCI groups (**Fig 5A**). Unlike the whole population of tSPNs, C_m_ was significantly decreased in NPY^+^ tSPNs (↓29.3%) (**Fig 5A_2_**). Numerical changes in R_in_ (↑18.0%) and τ_m_ (↓11.1%) were smaller than in the whole population and not statistically significant (**Fig 5A_3-4_**). Like the whole population, R_in_ was strongly correlated with τ_m_, but not C_m_ in both sham and SCI NPY^+^ tSPNs (**Fig 5B**).

**Table 2.** NPY^+^ tSPNs intrinsic properties in Sham and SCI groups. Intrinsic properties of NPY^+^ tSPNs. Values of basic properties of NPY^+^ tSPNs in sham and SCI groups. Data is presented as mean ± SD (number of observations). **RMP**, resting membrane potential; **R_in_**, input resistance; **τ_m_**, time constant; **C_m_**, capacitance; **I_rheo_**, rheobase; **T**, threshold; **Max dV/dt**, action potential maximum rising slope; **I_Na_**, sodium current; **I_A_**, A-type potassium current; **I_K_**, potassium delayed outward rectifier; **I_PIC_**, persistent inward current; **amp**, amplitude; ***f*_max_**, maximum instantaneous frequency; ***f*_sus_**, sustained firing frequency; **I**, current; *P < 0.05, ** P < 0.01. Unpaired student t-test.

|  | Variable | Sham | SCI | P value |
| --- | --- | --- | --- | --- |
| Passive properties | RMP, mV | -61.3±12.5 (12) | -63.9±9.4 (12) | 0.561 |
|  | R<sub>in</sub>, MΩ | 1858±1298 (11) | 2192±1840 (12) | 0.667 |
|  | τ<sub>m</sub>, ms | 149.9±90.0 (11) | 133.3±111.0 (12) | 0.389 |
|  | C<sub>m</sub>, pF | 92±37 (11) | 65±21 (12) | 0.045* |
| Spike threshold | I<sub>rheo</sub>, pA | 31.7±33.3 (11) | 16.6±11.1 (12) | 0.196 |
|  | Absolute T, mV | -42.3±7.5 (12) | -43.9±3.9 (12) | 0.516 |
|  | Relative T, mV | 31.5±13.2 (12) | 25.2±8.8 (12) | 0.180 |
| Spike waveform | Overshoot, mV | 31.0±14.5 (12) | 14.7±10.5 (12) | 0.005** |
|  | Amplitude, mV | 73.3±10.7 (12) | 58.7±12.9 (12) | 0.006** |
|  | Half width, ms | 4.5±1.8 (12) | 5.5±1.7 (12) | 0.046* |
|  | Max dV/dt, mV/ms | 71.0±25.2 (12) | 44.4±20.7 (12) | 0.010* |
| Voltage-gated currents | I<sub>Na</sub> amp, pA | 2751±1429 (11) | 1371±694 (12) | 0.007** |
|  | I<sub>A</sub> amp, pA | 1466±750 (12) | 708±316 (12) | 0.004** |
|  | I<sub>K</sub> amp, pA | 498±282 (11) | 322±154 (12) | 0.073 |
|  | I<sub>PIC</sub> amp, pA | 55.3±19.9 (8) | 38.8±26.5 (7) | 0.193 |
|  | I<sub>Na</sub> amp/C<sub>m</sub>, pA/pF | 32.9±14.8 (10) | 22.6±11.6 (12) | 0.080 |
|  | I<sub>A</sub> amp /C<sub>m</sub>, pA/pF | 16.6±6.0 (11) | 11.5±6.5 (12) | 0.027* |
|  | I<sub>K</sub> amp/C<sub>m</sub>, pA/pF | 5.6±3.0 (11) | 5.2±2.7 (12) | 0.667 |
|  | I<sub>PIC</sub> amp/C<sub>m</sub>, pA/pF | 0.08±0.11 (8) | 0.03±0.04 (7) | 0.397 |
| Repetitive Firing | f<sub>max</sub>-I slope, Hz/pA | 0.12±0.03 (12) | 0.17±0.05 (11) | 0.009** |
|  | f<sub>sus</sub>-I slope, Hz/pA | 0.05±0.03 (12) | 0.10±0.04 (10) | 0.004** |
|  | Spike Adaption Ratio | 1.75±0.42 (12) | 1.55±0.36 (10) | 0.251 |
|  | f<sub>max</sub> @ 60pA, Hz | 10.0±2.7 (10) | 13.4±3.1 (11) | 0.015* |
|  | f<sub>sus</sub> @ 60pA, Hz | 7.3±2.3 (10) | 9.7±2.4 (9) | 0.039* |

### (B) Changes in spike threshold properties in NPY^+^ tSPNs were similar to those in the whole population

Similar to the whole population, spike thresholds of NPY^+^ tSPNs were numerically reduced after SCI (↓1.8% in absolute T and ↓20% in relative T, **Fig 6A_2-3_**). The trend for numerical reductions in spike I_rheo_ (↓47.6%) and relative voltage threshold (↓20.0%) was more pronounced in NPY^+^ tSPNs after SCI (**Fig 6A**) than those seen in the whole population (↓31.2 and ↓13.1%, respectively). Consistent with whole population recordings, I_rheo_ was strongly correlated with input conductance (1/R_in_), but not with C_m_ or the inverse of τ_m_ (τ_m_^-1^) (**Fig 6B**). Detailed correlations between variables in NPY^+^ tSPNs are listed in **Sup Table 3**.

A smaller number of NPY^-^ tSPNs were recorded from the same NPY^TdTomato^ mice. Unlike NPY^+^ tSPNs, I_rheo_ was unchanged after SCI (**Table 3**), suggesting a different effect of SCI on excitability in functional subgroups.

**Table 3.** NPY^-^ tSPNs intrinsic properties in Sham and SCI groups. Intrinsic properties of NPY^-^ tSPNs. Values of basic properties of NPY^-^ tSPNs in sham and SCI groups. Data is presented as mean ± SD (number of observations). **RMP**, resting membrane potential; **R_in_**, input resistance; **τ_m_**, time constant; **C_m_**, capacitance; **I_rheo_**, rheobase; **T**, threshold; **Max dV/dt**, action potential maximum rising slope; **I_Na_**, sodium current; **I_A_**, A-type potassium current; **I_K_**, potassium delayed outward rectifier; **I_PIC_**, persistent inward current; **amp**, amplitude; ***f*_max_**, maximum instantaneous frequency; ***f*_sus_**, sustained firing frequency; **I**, current. Unpaired student t-test. *P < 0.05.

|  | Variable | Sham | SCI | P value |
| --- | --- | --- | --- | --- |
| Passive properties | RMP, mV | -53.0±4.6 (6) | -58.9±11.1 (7) | 0.239 |
|  | R<sub>in</sub>, MΩ | 791±384 (6) | 1418±1610 (7) | 1.000 |
|  | τ<sub>m</sub>, ms | 66.0±23.3 (5) | 95.5±99.4 (7) | 0.876 |
|  | C<sub>m</sub>, pF | 99±47.4 (5) | 80±30 (7) | 0.461 |
| Spike threshold | I<sub>rheo</sub>, pA | 39.6±19.1 (6) | 42.2±26.1 (6) | 0.848 |
|  | Absolute T, mV | -41.7±5.1 (6) | -46.9±2.7 (6) | 0.059 |
|  | Relative T, mV | 26.9±6.8 (6) | 22.0±1.5 (6) | 0.138 |
| Spike waveform | Overshoot, mV | 24.9±25.7 (6) | 23.6±15.9 (6) | 0.918 |
|  | Amplitude, mV | 66.6±26.5 (6) | 70.4±15.1 (6) | 0.768 |
|  | Half width, ms | 4.1±1.4 (6) | 3.9±1.2 (6) | 0.818 |
|  | Max dV/dt, mV/ms | 73.7±46.9 (6) | 68.8±27.4 (6) | 0.830 |
| Voltage-gated currents | I<sub>Na</sub> amp, pA | 1817±925 (3) | 1990±853 (5) | 0.805 |
|  | I<sub>A</sub> amp, pA | 1603±975 (5) | 1227±410 (7) | 0.452 |
|  | I<sub>K</sub> amp, pA | 639±208 (4) | 501±145 (7) | 0.297 |
|  | I<sub>PIC</sub> amp, pA | 59.1±36.8 (4) | 96.6±42.6 (3) | 0.266 |
|  | I<sub>Na</sub> amp/C<sub>m</sub>, pA/pF | 18.8±16.2 (3) | 31.3±21.5 (5) | 0.390 |
|  | I<sub>A</sub> amp /C<sub>m</sub>, pA/pF | 18.1±18.9 (4) | 17.3±8.8 (7) | 0.788 |
|  | I<sub>K</sub> amp/C<sub>m</sub>, pA/pF | 9.4±5.1 (3) | 7.4±4.7 (7) | 0.594 |
|  | I<sub>PIC</sub> amp/C<sub>m</sub>, pA/pF | 0.08±0.07 (4) | 0.18±0.21 (3) | 0.497 |
| Repetitive Firing | f<sub>max</sub>-I slope, Hz/pA | 0.11±0.05 (6) | 0.12±0.03 (5) | 0.585 |
|  | f<sub>sus</sub>-I slope, Hz/pA | 0.13±0.16 (6) | 0.07±0.01 (5) | 0.429 |
|  | Spike Adaption Ratio | 1.52±0.30 (6) | 1.56±0.22 (5) | 0.816 |
|  | f<sub>max</sub> @ 60pA, Hz | 9.8±4.0 (4) | 8.2±2.0 (4) | 0.516 |
|  | f<sub>sus</sub> @ 60pA, Hz | 6.3±2.1 (4) | 6.9±2.1 (4) | 0.486 |

### (C) Higher reductions in I_Na_ and I_A_ in NPY^+^ tSPNs contribute to changes in action potential waveform after SCI

Unlike the whole population, action potentials were significantly broader (half width ↑22.2%) and smaller (amplitude ↓19.9%) in NPY^+^ tSPNs after SCI (**Fig 6C**) and were associated with significantly larger reductions in I_Na_ (↓50.2%) and I_A_ (↓51.7%) amplitudes (**Fig 6D**) compared to the whole population (↓31.1% and ↓33.5%, respectively). I_Na_ current density numerically decreased 31.3% and I_A_ current density significantly decreased 30.7% in NPY^+^ tSPNs, compared to the numerical decreases of 14.8% and 19.6%, respectively, in the whole population (**Table 2**). Collectively these results demonstrate that, in NPY^+^ tSPNs, SCI leads to reductions in cell size and channel density, resulting in smaller and broader spikes. Consistent with the whole population, I_Na_ and I_A_ magnitudes in NPY^+^ tSPNs were similarly correlated in sham and SCI populations (**Fig 6E**). Although there were significant correlations between spike waveform and I_Na_ and I_A_ amplitudes, they were less significant after SCI, suggesting that additional currents contribute to spike shape after SCI [e.g. delayed outward rectifier^44^] (**Fig 6F, Sup Table 4**).

Contrary to NPY^+^ tSPNs, in NPY^-^ tSPNs, I_Na_ amplitude was unchanged, while I_A_ amplitude numerically decreased 23.5% after SCI (**Table 3**), providing further evidence that SCI effects are not uniform among neuronal subpopulations.

### (D) NPY^+^ tSPNs had increased output gain after SCI

Repetitive firing properties were assessed with *f*-I curves in NPY^+^ tSPNs. Compared to the broad distribution seen in the whole population of tSPNs recordings, *f*-I curves in SCI mice clustered toward higher excitability levels, with *f*_max_-I and *f*_sus_-I curve slopes being significantly increased compared to sham (**Fig 7A_1-3_, 7B_1-3_**). When given a slight suprathreshold injected current (60pA), both *f*_max_ and *f*_sus_ were significantly higher after SCI (**Fig 7A_5_, 7B_5_**). Moderate correlations between *f*_max_-I curve slopes and R_in_ and I_rheo_ were similar with those observed in the whole population (**Fig 7C**).

To further verify the enhanced output gain after SCI in NPY^+^ tSPNs, we used a computational model and simulated *f*_max_-I and *f*_sus_-I curves using mean value ratio changes in C_m_, R_in_, I_NA_, I_A_ and I_K_ between sham and SCI groups comparable to those given in **Table 2**. The model showed a leftward shift and an increase in *f*-I curve slope using SCI parameters compared to sham (**Fig 7A_4_, 7B_4_**), where the adjusted parameters collectively contributed to the modeled outcome, rather than any single parameter acting as the primary driver (as detailed later). These results suggest that changes in these membrane properties can account for the observed increased excitability and output gain in NPY^+^ tSPNs following SCI.

### (E) Persistent inward current contributions to repetitive firing in tSPNs

Voltage-gated Na^+^ and Ca^2+^ conductances can generate persistent inward currents (PICs) that enhance excitability and support sustained firing^45^. As all tSPNs were capable of sustained firing, voltage ramps were delivered to assess their PICs properties. PICs were observed in ∼2/3 tSPNs regardless of population (whole, NPY^+^, NPY^-^) in both sham (68%, 67%, 67%) and SCI mice (77%, 70%, 60%), respectively (P<0.5-0.9). PIC amplitudes were comparable between sham and SCI tSPNs in the whole population (**Table 1**). That PIC amplitudes were ∼double the I_rheo_ currents required to initiate spiking (**Table 1**) supports a PIC contribution to sustained firing. PICs were numerically smaller after SCI in NPY^+^ (↓30%) but larger in NPY^-^ (↑63.5%) tSPNs (**Table 2-3, Sup Fig 2A**). PIC thresholds were also comparable between sham and SCI in the whole population (-46.4±10.1mV [n=13] and-50.8±11.8mV [n=23], respectively), but were significantly leftward-shifted after SCI in NPY^+^ tSPNs (-42.9±7.1mV [n=8] in sham vs - 56.8±15.8mV [n=7] in SCI; P=0.04) (**Sup Fig 2B**). The leftward shift in PIC threshold in the NPY^+^ population after SCI is consistent with the observed significant increase in *f*-I slope. That mean PIC threshold was ∼17 mV more negative than absolute T in the NPY^+^ SCI population would support a PIC contribution to spike recruitment (**Sup Table 3**). In contrast to NPY^+^ tSPNs, a trend toward an opposite right-ward shift was seen in NPY^-^ tSPNs PICs threshold (average - 52.1 ± 14.3mV of 4 tSPNs in sham, and-39.0 ± 3.3mV of 3 tSPNs in SCI, respectively) (**Sup Fig 2B**).

The L-type Ca^2+^ channel blocker nifedipine fully blocked PICs in 4/5 tSPNs, while subsequent Na^+^ channel blocker TTX blocked the PIC in the remaining tSPN (not shown; **Sup Fig 2C-D**). These data demonstrate that L-type Ca^2+^ channels contribute to tSPN neuronal excitability and spike frequency regulation.

### (F) Computational model assessment of channel conductances contributing to firing properties

We systemically manipulated variables in the optimized computational model to explore factors capable of changing NPY^+^ tSPNs *f*-I curves. A model NPY^+^ tSPN was simulated to mimic the average *f*-I curves observed in sham NPY^+^ tSPNs recordings. We then decreased (indicated by ↓) or increased (indicated by ↑) parameters consistent with experimental changes of variables observed in NPY^+^ tSPNs after SCI: ↓I_Na_, ↓I_A_, ↓I_K_, ↓C_m_, and ↑R_in_ (implemented by ↓G_leak_). I_K_ was correlated with *f*-I slopes after SCI (**Sup Table 3**), and ↓I_K_ was the most effective parameter change for increasing model *f*-I slopes (*f*_max_-I and *f*_sus_-I). Comparison of effects of other parameters and conductances on *f*-I curves is provided in (**Sup Table 4**).

## Synaptic input

### (A) tSPNs have increased spontaneous synaptic activity after SCI

Almost all recorded tSPNs received some degree of spontaneous synaptic activity (**Table 4**). Measured as spontaneous excitatory postsynaptic currents (sEPSC), these events were resistant to TTX (**Fig 8A_1-2_**), demonstrating that these are spike-independent quantal synaptic events. sEPSC frequency was significantly increased in both the whole population and NPY^+^ tSPNs after SCI (2.3 times and 3 times, respectively) (**Fig 8B**). Changes were unrelated to variation in passive membrane properties (**Sup Table 5**). There were no changes in sEPSC frequency in the NPY^-^ population (**Fig 8B**). sEPSC amplitudes were not significantly different between sham and SCI populations (**Table 4; Sup Fig 3A**). sEPSC decay time (τ_decay_) was unchanged overall and in NPY^+^ tSPNs, but significantly reduced in NPY^-^ tSPNs (31.7%). (**Table 4; Sup Fig 3B**).

**Table 4.** tSPNs received sEPSC properties in sham and SCI groups. tSPNs received sEPSC properties in sham and SCI groups. Incidence is p resented with number of tSPNs that received sEPSC/total number of recorded tSPN in the category. Other variables are presented as mean ± SD (number of observations). ***f***, frequency; **τ_decay_**, time constant of sEPSC decay phase. Unpaired student t-test. *P < 0.05, ** P < 0.01.

|  | Whole population tSPN |  |  | NPY<sup>+</sup> tSPN |  |  | NPY<sup>-</sup> tSPN |  |  |
| --- | --- | --- | --- | --- | --- | --- | --- | --- | --- |
|  | Sham | SCI | P value | Sham | SCI | P value | Sham | SCI | P value |
| Incidence | 16/17 | 35/35 |  | 9/10 | 11/11 |  | 6/6 | 7/7 |  |
| *f*, Hz | 1.9±2.2 (17) | 4.3±3.8 (35) | 0.007** | 1.4±1.7 (10) | 4.2±4.7 (11) | 0.038* | 2.0±2.4 (6) | 2.9±1.3 (7) | 0.472 |
| Amplitude, pA | 20.0±11.3 (16) | 18.0±11.7 (35) | 0.641 | 20.9±13.7 (9) | 14.3±4.6 (11) | 0.382 | 19.5±8.6 (6) | 14.6±2.7 (7) | 0.174 |
| $\tau_{\text{decay}}$, ms | 9.3±4.6 (16) | 7.4±3.1 (35) | 0.200 | 9.3±5.7 (9) | 7.0±3.8 (11) | 0.409 | 10.1±3.2 (6) | 6.9±2.0 (7) | 0.046* |

Physiologically, tSPN recruitment largely depends on preganglionic excitatory synaptic drive, determined by excitatory postsynaptic potential (EPSP) amplitude and synaptic summation relative to the voltage threshold for spiking. In current clamp, spontaneous EPSPs (sEPSPs) were capable of recruiting spikes (**Fig 8C_1_**). The proportion of whole population tSPNs exhibiting spontaneous spikes was numerically higher after SCI compared to sham (22.2% vs. 10.5%, respectively; P=0.28) (**Fig 8C_2_**).

### (B) Computational modeling supports larger EPSPs and increased recruitment after SCI

To explore the overall impact of changes in tSPN intrinsic properties and frequency of spontaneous synaptic activity on tSPN recruitment after SCI, we delivered synaptic currents to model neurons that represented the intrinsic properties of tSPNs from sham and SCI experimental groups. A stronger synaptic current amplitude was used in the model than obtained experimentally because the single-compartment model introduces strong shunting interactions between synaptic inputs compared to that seen physiologically^46^. In response to the same synaptic current, the amplitude of the unitary EPSP was greater in the SCI model neuron compared to the sham model, suggesting that SCI-induced changes in the intrinsic properties of tSPNs increase tSPN excitability and enhance the efficacy of preganglionic input (**Fig 8D_1_**). This increased excitability in the SCI model neuron contributed to a higher frequency of spiking compared to the sham model (**Fig 8D_2_**). Although a single EPSP was insufficient to trigger action potentials, EPSPs could summate to produce spiking with EPSCs at frequencies observed experimentally (**Fig 8D_2_**). The increased sEPSC frequency after SCI resulted in a higher firing rate in the SCI model neuron (**Fig 8D_2_**), indicating that more frequent spontaneous synaptic input enhances tSPNs recruitment.

## Population recordings from the intact thoracic chain

### (A) Spontaneous synaptic activity can recruit spiking in tSPN axons in interganglionic nerves

Recordings from individual neurons demonstrated that ongoing quantal spontaneous synaptic activity can recruit spiking in tSPNs. To examine this further, we sought to demonstrate that spontaneous synaptic activity could recruit spiking in tSPN axons in an intact T4-T12 *ex vivo* sympathetic chain preparation (**Fig 8E**). Spiking activity in axons adjacent to rostral T4 and caudal T12 ganglia were recorded using tight suction electrodes on interganglionic nerves (IGNs) before and after block of synaptic transmission with hexamethonium (Hex) from both naive-sham (n=9) and SCI (n=15) mice. Few tSPN axons are expected to be found in IGN recordings as most exit through spinal nerves at the same segment ^47^. Even so, Hex-sensitive spontaneous activity was seen in 47% (22/47) of IGN recordings (9/17 naive-sham and 13/30 SCI IGNs; p=0.53). Thus, spontaneous synaptic activity was sufficient to recruit axonal spiking in tSPN axons in the intact chain.

### (B) After SCI, preganglionic axonal recruitment was reduced in caudally-projecting axons while rostrally projecting axons may lead to response amplification

The *ex vivo* intact thoracic sympathetic chain also contained connected ventral roots (**Fig 8E**). We selectively recruited T10 spinal preganglionic axons via supramaximal T10 ventral root stimulation while recording evoked compound action potential (CAP) responses in the IGN from caudally-projecting preganglionic axons at T12 and rostrally-projecting axons at T4 (propagation across 3 vs. 7 ganglia, respectively; **Fig 8E**; **Table 5**). Evoked CAPs comprise directly-activated preganglionic and synaptically-evoked tSPN components. T12 vs. T4 CAPs were larger by 427% in sham-naive (p=0.03) and 284% in SCI (p=0.07) (**Table 5**). After SCI, evoked CAPs trended toward reduction at T12 (↓50%; p=0.13) but not T4 (↓24%; p=0.54).

**Table 5.** Evoked compound action potentials in sham/naive and SCI groups. Evoked compound action potentials in sham/naive and SCI groups. Baseline noise subtracted rectified integral values of evoked responses at rostral and caudal thoracic ganglia. Data is presented as mean ± SD (number of observations). **CAP**, compound action potential; **hex**, hexamethonium. Unpaired student t-test. *P < 0.05.

|  | Ganglia | Sham/Naive | SCI | P value |
| --- | --- | --- | --- | --- |
| Total Evoked Response | T4 | 175.7±144.1 (6) | 133.2±90.4 (9) | 0.54 |
|  | T12 | 750.0±537.3 (6) | 377.7±368.6 (9) | 0.13 |
| Preganglionic CAP<br>(response after hex) | T4 | 119.0±94.3 (6) | 94.3±81.4 (9) | 0.41 |
|  | T12 | 440.8±300.1 (6) | 191.7±147.0 (9) | 0.05* |
| Postganglionic CAP<br>(Total - response after hex) | T4 | 56.7±54.1 (6) | 38.9±40.4 (9) | 0.49 |
|  | T12 | 309.2±277.4 (6) | 186.0±249.4 (9) | 0.11 |
| Postganglionic fraction<br>(Post-/Preganglionic) | T4 | 0.43±0.30 (6) | 1.69±2.80 (9) | 0.21 |
|  | T12 | 0.82±0.47 (6) | 1.14±0.77 (9) | 0.39 |
| Population synaptic<br>amplitude (μV) | T4 | 32.9±34.2 (6) | 25.8±20.8 (9) | 0.62 |
|  | T12 | 161.8±119.3 (6) | 63.7±56.1 (9) | 0.05* |

The preganglionic component of the CAP, obtained following synaptic block of tSPN activity with Hex, was significantly reduced after SCI at T12 (↓57%; p=0.05) but not at T4 (↓21%; p=0.41), indicating a preferential loss of caudally-directed axon collaterals (**Fig 8F**; **Table 5**). As supramaximal stimuli were applied, it is unlikely that changes in recruitment are due to changes in the electrical excitability of these axons.

The postganglionic (tSPN) component of the CAP also trended toward preferential reduction after SCI at T12 (↓40%; p=0.11). Indeed, the low-pass filtered subthreshold population EPSP responses were significantly smaller after SCI at T12 (↓61%; p=0.05) (**Fig 8G**; **Table 5**).

That Hex significantly reduced CAP amplitude of all evoked volleys (sham-naive: T4 [p=0.05]; T12 [p=0.031] and SCI T4 [p=0.02]; T12 [p=0.004]), demonstrates that a single synchronous stimulus-evoked preganglionic volley was capable of significant suprathreshold synaptic recruitment of tSPNs in ganglia many segments away. Using the postganglionic/preganglionic volley fraction to compare relative tSPN axon recruitment, there is ∼400% greater recruitment in SCI vs sham-naive mice at T4 (1.69 vs. 0.43, respectively) compared to ∼40% at T12 (1.14 vs. 0.84, respectively) (**Table 5**). Though differences were not significant, output amplification in rostrally-projecting axons after SCI was most probable. The possibility of differential plasticity between caudal and rostral sympathetic ganglia requires further investigation.

## Discussion

### SCI-induced plasticity in the overall tSPNs populations may be masked by underlying heterogeneity

Thoracic SPNs are functionally heterogenous, comprising seven molecularly distinct populations likely associated with their specific target organs^28,48^. Given the critical role of tSPNs in maintaining organismal homeostasis, we comprehensively evaluated whether they undergo compensatory increases in cellular excitability following the loss of descending sympathetic drive several weeks post-high thoracic SCI. Broadly speaking, surprisingly few statistically significant SCI-induced changes were seen in whole population tSPN passive and active membrane properties, suggesting remarkable resilience to lost central drive. However, these negative findings must be interpreted cautiously due to the enormous range in population cellular excitability. tSPN recruitment threshold occurred over a >10-fold range in required current (I_rheo_) and was driven by a largely ohmic >10-fold range in cell resistance (R_in_). While voltage-gated current densities (I_Na_, I_A_, I_K_, I_PIC_), spike waveforms and firing properties showed no significant changes after SCI, a numerical decrease in I_rheo_ suggested a trend toward increased excitability, potentially facilitating recruitment by preganglionic inputs.

Large variability was observed in nearly all assessed electrophysiological parameters, albeit with narrower range in a functional subpopulation (see below). Substantial variability within a population greatly reduces statistical power (1-β) for group comparisons. Much larger estimated sample sizes to achieve 1-β ≥ 0.8 are impractical given the technical challenges of whole-cell patching in sympathetic ganglia and time-intensive high thoracic spinalizations. Still, consistent numerical trends and many significant changes, combined with computational modeling, provide a comprehensive understanding of these neurons and their plasticity after SCI as further elaborated below.

tSPN morphology was also highly variable in soma size and dendritic complexity. Nonetheless, dendritic complexity was significantly reduced after SCI, potentially representing overall cell shrinkage associated with numeric reduction in cell capacitance (↓C_m_). Numeric increases in R_in_ and τ_m_ align with these changes and support an increased capacity for synaptic integration.

### Considerations on apparent resilience of membrane properties after SCI

High thoracic SCI produces profound depression of autonomic activity due to loss of central drive, leaving minimal ongoing SPN activity to somatic structures such as skin and muscle, which are predominantly innervated by paravertebral SPNs^49,50^. Given this, homeostatic plasticity-induced increases in SPN excitability may be expected^51^, but possibly blunted by ongoing clinically-comparable^53^ episodes of exaggerated sympathetic drive associated with AD^26,52^. However, across the whole population, tSPN excitability appeared unchanged after SCI suggesting that tSPNs are inherently resilient to SCI-associated changes in preganglionic drive. For example, blocking synaptic input in chick embryo SPNs did not observe the expected excitability increases^54^. Conversely, overall changes may mask subpopulation-specific plasticity as seen with reductions in I_rheo_ and I_NA_ in NPY^+^ tSPNs but not in NPY^-^ tSPNs.

### Putative vasomotor NPY^+^ tSPNs exhibit enhanced intrinsic excitability after SCI

Two of the five TH^+^ tSPN subpopulations express NPY and comprise ∼50% of all tSPNs^28^. Approximately 80% of NPY^+^ tSPNs (the NA3 group^28^) are smaller in diameter and presumed to innervate vasculature^16,28,30^. Fluorescently-identified NPY^+^ tSPNs had a 33.5% lower C_m_ than the whole population, consistent with preferential sampling from the NA3 vasomotor group.

NPY^+^ tSPNs also showed reduced variability in membrane properties regardless of injury when compared to the whole population. For instance, C_m_ varied over a >7-fold range in the whole population, but only >3-fold in NPY^+^ tSPNs, increasing the likelihood of detecting statistically significant changes. Indeed, after SCI, NPY^+^ tSPNs were significantly smaller in cell size (↓C_m_) with more significant reductions in I_Na_ and I_A_, altering spike waveforms. Significantly increased firing frequency and *f*-I slopes after SCI indicate enhanced output gain. Persistent inward currents (PICs) were common, and a leftward shift in PIC threshold after SCI may contribute to the increased excitability of NPY^+^ tSPNs. Computational modeling indicated other conductances, particularly delayed rectifiers (↓I_K_), also shift *f*-I curves.

PICs are generated by Na_V_ and L-type Ca^2+^ channels ^34,55^. Na_V_1.7 is the predominant and most expressed Na_V_ channel in tSPNs^28^. Its slow recovery from fast inactivation could contribute to the generation of PICs^56,57^. Nonetheless, nifedipine-sensitivity identified Ca^2+^ channels as predominantly responsible for PICs in most NPY^+^ tSPNs. That tSPNs have limited transcript expression of Cacna1d (Ca_V_1.3) L-type Ca^2+^ channels^28^ aligns with small amplitude PICs generated by their activation.

Consistent with previous findings^19,48^, tSPN recruitment in both sham and SCI groups was strongly driven by R_in_. This recruitment principle inherently favors the activation of vasculature-innervating NPY^+^ tSPNs, which exhibit higher R_in_ than NPY^-^ tSPNs. The observed preferential increases in spontaneous synaptic activity in NPY^+^ tSPNs after SCI further favors their recruitment (see below). Conversely, NPY^-^ tSPNs showed increased I_rheo_ and a decreased *f*-I slopes trending toward reduced excitability after SCI. These findings suggest that distinct functional subpopulations of tSPNs may undergo divergent forms of plasticity following SCI.

### Changes in spontaneous synaptic activity support increased tSPN recruitment

Spontaneous synaptic activity occurred in nearly all tSPNs, consistent with previous findings in guinea-pig^17^. Assuming SCI dramatically reduces preganglionic drive, one may expect homeostatic synaptic scaling in sEPSC amplitude^58^. This was not seen, suggesting synaptic scaling may not be a homeostatic mechanism occurring in tSPNs, perhaps due to their low synapse number^59^ and presence of suprathreshold synapses^15^. Alternatively, the occasional epochs of strong preganglionic drive may counteract overall reductions in drive seen.

There was substantial variability in quantal sEPSC frequency across individual neurons. Following SCI, both the whole population and NPY^+^ (but not NPY^-^) tSPNs exhibited a significant sEPSC frequency increase. Observed increases in spontaneous frequency after SCI are indicative of more synapses and/or increased release probability^60^. The increased sEPSC frequency after SCI is consistent with a homeostatic response to reduction in synaptic transmission as previously seen in the adult mouse superior cervical ganglia following antibody-mediated impairment of sympathetic ganglionic transmission^61^. sEPSC frequency increases after SCI are consistent with a reduction in ongoing evoked responses, since evoked responses reduce spontaneous frequency^62^. This is consistent with the expectations of reduced central drive following SCI. An increased frequency may also be due to increases in synapse number arising from preganglionic axon collateral sprouting. Denervation of sympathetic ganglia leads to extensive sprouting of residual preganglionic axons within the ganglia^63,64^ and SCI is expected to lead to a partial denervation due to destruction of preganglionics at the injury site. However, given that we observed morphological and electrophysiological evidence of reduction in tSPN size and dendritic complexity, it is likely there is limited synaptic sprouting.

Consistent with our previous study in tSPNs from naive mice^19^, whole-cell recordings revealed that sEPSP summation generated spontaneous firing, occurring in 10% of sham and 22% of SCI tSPNs. Computational modeling of synaptic actions in NPY^+^ tSPNs demonstrated that spontaneous inputs increase spike recruitment post-SCI. Population recordings from the intact thoracic chain confirmed that spontaneous synaptic activity can recruit spiking in tSPN axons in interganglionic nerves at rostral (T4/5) and caudal (T11/12) locations with intact multisegmental connections. Unlike sEPSC recordings in tSPNs, where recruitment numerically doubled, there were no differences in incidence from tSPN axons recruited in the interganglionic nerve (IGN). However, only a small fraction of tSPNs project axons through the IGN^47^, and these may represent a specific subpopulation with low spontaneous synaptic activity.

Regardless of mechanism, increased sEPSC frequency would be expected to increase evoked response amplitudes during preganglionic recruitment (e.g. during AD). This is because spontaneous and evoked synaptic responses arise from the same readily releasable pool, and the pool size strongly correlates with spontaneous frequency and evoked synaptic amplitude ^65,66^. Our studies could not directly test this. Cellular recordings were obtained predominantly from T3-5 ganglia, which receive most input from preganglionics at the same or nearby spinal segments^67^. Unfortunately, our assessment of population evoked responses at the T4 ganglia was undertaken in the intact thoracic paravertebral chain following stimulation of preganglionic axons from a distant segment (T10) expected to provide limited synaptic drive. While the preganglionic volley may recruit a larger fraction of postganglionic axons (∼400% numerical increase), estimated population synaptic response was unchanged.

## Conclusions

In sum, while tSPN membrane properties were highly heterogenous, with some varying over a >10-fold range, SCI-induced changes were consistent with increased cellular and synaptic excitability. These findings support tSPNs as an additional site of excitability amplification in the sympathetic circuit after SCI. While high-thoracic SCI reduces preganglionic drive to tSPNs, causing insufficient vasoconstriction and impaired hemodynamic regulation, increased NPY^+^ tSPNs excitability may partially compensate by restoring basal vascular tone, critical for maintaining blood pressure. However, intrinsic amplification would also support exaggerated responses generated by afferent-driven preganglionic input, contributing to dysregulated vasoconstriction and the development of AD.

Enhanced spontaneous activity would promote CNS-independent actions and may serve as a compensatory mechanism to partially restore basal sympathetic drive to effector organs including NPY^+^ vasoconstrictors. Both cellular and population recordings of tSPN axons in interganglionic nerves in the intact chain show the capacity for recruiting spontaneous spiking. Further, computational modeling supports larger EPSPs and increased recruitment in NPY^+^ tSPNs after SCI.

## References

1. Teasell, R.W., Arnold, J.M., Krassioukov, A. & Delaney, G.A. Cardiovascular consequences of loss of supraspinal control of the sympathetic nervous system after spinal cord injury. Arch Phys Med Rehabil 81, 506–516 (2000).

2. Krassioukov, A., Warburton, D.E., Teasell, R., Eng, J.J. & Spinal Cord Injury Rehabilitation Evidence Research, T. A systematic review of the management of autonomic dysreflexia after spinal cord injury. Arch Phys Med Rehabil 90, 682–695 (2009).

3. White, A.R. & Holmes, G.M. Anatomical and Functional Changes to the Colonic Neuromuscular Compartment after Experimental Spinal Cord Injury. J Neurotrauma 35, 1079–1090 (2018).

4. Qi, Z., Middleton, J.W. & Malcolm, A. Bowel Dysfunction in Spinal Cord Injury. Curr Gastroenterol Rep 20, 47 (2018).

5. Wan, D. & Krassioukov, A.V. Life-threatening outcomes associated with autonomic dysreflexia: a clinical review. J Spinal Cord Med 37, 2–10 (2014).

6. Eldahan, K.C. & Rabchevsky, A.G. Autonomic dysreflexia after spinal cord injury: Systemic pathophysiology and methods of management. Auton Neurosci 209, 59–70 (2018).

7. Rabchevsky, A.G. Segmental organization of spinal reflexes mediating autonomic dysreflexia after spinal cord injury. Prog Brain Res 152, 265–274 (2006).

8. Weaver, L.C., Marsh, D.R., Gris, D., Brown, A. & Dekaban, G.A. Autonomic dysreflexia after spinal cord injury: central mechanisms and strategies for prevention. Prog Brain Res 152, 245–263 (2006).

9. Ueno, M., Ueno-Nakamura, Y., Niehaus, J., Popovich, P.G. & Yoshida, Y. Silencing spinal interneurons inhibits immune suppressive autonomic reflexes caused by spinal cord injury. Nature neuroscience 19, 784–787 (2016).

10. Brock, J.A., Yeoh, M. & McLachlan, E.M. Enhanced neurally evoked responses and inhibition of norepinephrine reuptake in rat mesenteric arteries after spinal transection. Am J Physiol Heart Circ Physiol 290, H398–405 (2006).

11. Yeoh, M., McLachlan, E.M. & Brock, J.A. Tail arteries from chronically spinalized rats have potentiated responses to nerve stimulation in vitro. J Physiol 556, 545–555 (2004).

12. Lujan, H.L., Tonson, A., Wiseman, R.W. & DiCarlo, S.E. Chronic, complete cervical(6-7) cord transection: distinct autonomic and cardiac deficits. J Appl Physiol (1985) 124, 1471–1482 (2018).

13. Jänig, W. The integrative action of the autonomic nervous system: neurobiology of homeostasis, (Cambridge University Press, Cambridge, United Kingdom; New York, NY, 2022).

14. Purves, D., Rubin, E., Snider, W.D. & Lichtman, J. Relation of animal size to convergence, divergence, and neuronal number in peripheral sympathetic pathways. J Neurosci 6, 158–163 (1986).

15. McLachlan, E.M. Transmission of signals through sympathetic ganglia--modulation, integration or simply distribution? Acta physiologica Scandinavica 177, 227–235 (2003).

16. Jobling, P. & Gibbins, I.L. Electrophysiological and morphological diversity of mouse sympathetic neurons. Journal of neurophysiology 82, 2747–2764 (1999).

17. Blackman, J.G. & Purves, R.D. Intracellular recordings from ganglia of the thoracic sympathetic chain of the guinea-pig. J Physiol 203, 173–198 (1969).

18. Lichtman, J.W., Purves, D. & Yip, J.W. Innervation of sympathetic neurones in the guinea-pig thoracic chain. J Physiol. 298, 285–299. (1980).

19. McKinnon, M.L., et al. Dramatically Amplified Thoracic Sympathetic Postganglionic Excitability and Integrative Capacity Revealed with Whole-Cell Patch-Clamp Recordings. eNeuro 6(2019).

20. Springer, M.G., Kullmann, P.H. & Horn, J.P. Virtual leak channels modulate firing dynamics and synaptic integration in rat sympathetic neurons: implications for ganglionic transmission in vivo. The Journal of physiology 593, 803–823 (2015).

21. Karila, P. & Horn, J.P. Secondary nicotinic synapses on sympathetic B neurons and their putative role in ganglionic amplification of activity. J Neurosci 20, 908–918 (2000).

22. Rimmer, K. & Horn, J.P. Weak and straddling secondary nicotinic synapses can drive firing in rat sympathetic neurons and thereby contribute to ganglionic amplification. Frontiers in neurology 1, 130 (2010).

23. Janig, W., Krauspe, R. & Wiedersatz, G. Activation of postganglionic neurones via non-nicotinic synaptic mechanisms by stimulation of thin preganglionic axons. Pflugers Archiv: European journal of physiology 401, 318–320 (1984).

24. Jacob, J.E., Pniak, A., Weaver, L.C. & Brown, A. Autonomic dysreflexia in a mouse model of spinal cord injury. Neuroscience 108, 687–693 (2001).

25. Osei-Owusu, P., Collyer, E., Dahlen, S.A., Adams, R.E. & Tom, V.J. Maladaptation of renal hemodynamics contributes to kidney dysfunction resulting from thoracic spinal cord injury in mice. Am J Physiol Renal Physiol 323, F120–F140 (2022).

26. Brennan, F.H., et al. Microglia promote maladaptive plasticity in autonomic circuitry after spinal cord injury in mice. Sci Transl Med 16, eadi3259 (2024).

27. Savitt, J.M., Jang, S.S., Mu, W., Dawson, V.L. & Dawson, T.M. Bcl-x is required for proper development of the mouse substantia nigra. The Journal of neuroscience: the official journal of the Society for Neuroscience 25, 6721–6728 (2005).

28. Furlan, A., et al. Visceral motor neuron diversity delineates a cellular basis for nipple-and pilo-erection muscle control. Nature neuroscience 19, 1331–1340 (2016).

29. Elfvin, L.G., Lindh, B. & Hokfelt, T. The chemical neuroanatomy of sympathetic ganglia. Annual review of neuroscience 16, 471–507 (1993).

30. Gibbins, I.L. Vasomotor, pilomotor and secretomotor neurons distinguished by size and neuropeptide content in superior cervical ganglia of mice. Journal of the autonomic nervous system 34, 171–183 (1991).

31. Halder, M., McKinnon, M.L., Li, Y., Wenner, P. & Hochman, S. Isolation and Electrophysiology of Murine Sympathetic Postganglionic Neurons in the Thoracic Paravertebral Ganglia. Bio Protoc 11, e4189 (2021).

32. Golowasch, J., et al. Membrane capacitance measurements revisited: dependence of capacitance value on measurement method in nonisopotential neurons. Journal of neurophysiology 102, 2161–2175 (2009).

33. Dougherty, K.J. & Hochman, S. Spinal cord injury causes plasticity in a subpopulation of lamina I GABAergic interneurons. J Neurophysiol 100, 212–223 (2008).

34. Li, Y., Gorassini, M.A. & Bennett, D.J. Role of persistent sodium and calcium currents in motoneuron firing and spasticity in chronic spinal rats. Journal of neurophysiology 91, 767–783 (2004).

35. Schild, J.H., et al. A-and C-type rat nodose sensory neurons: model interpretations of dynamic discharge characteristics. J Neurophysiol 71, 2338–2358 (1994).

36. Günay, C. & Prinz, A.A. An offline correction method for uncompensated series resistance and capacitance artifacts from whole-cell patch clamp recordings of small cells. BMC Neuroscience 12, P259 (2011).

37. Dusterwald, K.M., et al. Biophysical models reveal the relative importance of transporter proteins and impermeant anions in chloride homeostasis. Elife 7(2018).

38. Jarve, A., et al. Distinct roles of angiotensin receptors in autonomic dysreflexia following high-level spinal cord injury in mice. Exp Neurol 311, 173–181 (2019).

39. McKinnon, M.L., et al. Dramatically Amplified Thoracic Sympathetic Postganglionic Excitability and Integrative Capacity Revealed with Whole-Cell Patch-Clamp Recordings. eNeuro 6, ENEURO.0433-0418.2019 (2019).

40. Gustafsson, B. & Pinter, M.J. An investigation of threshold properties among cat spinal alpha-motoneurones. The Journal of physiology 357, 453–483 (1984).

41. Belluzzi, O., Sacchi, O. & Wanke, E. A fast transient outward current in the rat sympathetic neurone studied under voltage-clamp conditions. J Physiol 358, 91–108 (1985).

42. Zimmerman, A. & Hochman, S. Heterogeneity of membrane properties in sympathetic preganglionic neurons of neonatal mice: evidence of four subpopulations in the intermediolateral nucleus. Journal of neurophysiology 103, 490–498 (2010).

43. Morris, J.L. Selective constriction of small cutaneous arteries by NPY matches distribution of NPY in sympathetic axons. Regulatory peptides 49, 225–236 (1994).

44. Baranauskas, G. Ionic channel function in action potential generation: current perspective. Mol Neurobiol 35, 129–150 (2007).

45. Heckmann, C.J., Gorassini, M.A. & Bennett, D.J. Persistent inward currents in motoneuron dendrites: implications for motor output. Muscle & nerve 31, 135–156 (2005).

46. Nelson, M.E. A Mechanism for Neuronal Gain Control by Descending Pathways. Neural Computation 6, 242–254 (1994).

47. Jänig, W. The integrative action of the autonomic nervous system: neurobiology of homeostasis, (Cambridge University Press, Cambridge, 2008).

48. Hochman, S. & McCrea, D.A. Effects of chronic spinalization on ankle extensor motoneurons. II. Motoneuron electrical properties. Journal of neurophysiology 71, 1468–1479 (1994).

49. Stjernberg, L., Blumberg, H. & Wallin, B.G. Sympathetic activity in man after spinal cord injury. Outflow to muscle below the lesion. Brain 109 ( Pt 4), 695–715 (1986).

50. Wallin, B.G. & Stjernberg, L. Sympathetic activity in man after spinal cord injury. Outflow to skin below the lesion. Brain 107 ( Pt 1), 183–198 (1984).

51. Wen, W. & Turrigiano, G.G. Keeping Your Brain in Balance: Homeostatic Regulation of Network Function. Annu Rev Neurosci 47, 41–61 (2024).

52. Zhang, Y., et al. Autonomic dysreflexia causes chronic immune suppression after spinal cord injury. The Journal of neuroscience: the official journal of the Society for Neuroscience 33, 12970–12981 (2013).

53. Hubli, M., Gee, C.M. & Krassioukov, A.V. Refined assessment of blood pressure instability after spinal cord injury. Am J Hypertens 28, 173–181 (2015).

54. Ratliff, A., Pekala, D. & Wenner, P. Plasticity in Preganglionic and Postganglionic Neurons of the Sympathetic Nervous System during Embryonic Development. eNeuro 10(2023).

55. Carlin, K.P., Jones, K.E., Jiang, Z., Jordan, L.M. & Brownstone, R.M. Dendritic L-type calcium currents in mouse spinal motoneurons: implications for bistability. The European journal of neuroscience 12, 1635–1646 (2000).

56. Cummins, T.R., Howe, J.R. & Waxman, S.G. Slow closed-state inactivation: a novel mechanism underlying ramp currents in cells expressing the hNE/PN1 sodium channel. J Neurosci 18, 9607–9619 (1998).

57. Cummins, T.R., Sheets, P.L. & Waxman, S.G. The roles of sodium channels in nociception: Implications for mechanisms of pain. Pain 131, 243–257 (2007).

58. Turrigiano, G.G. & Nelson, S.B. Homeostatic plasticity in the developing nervous system. Nature reviews. Neuroscience 5, 97–107 (2004).

59. Gibbins, I.L., Jobling, P., Messenger, J.P., Teo, E.H. & Morris, J.L. Neuronal morphology and the synaptic organisation of sympathetic ganglia. Journal of the autonomic nervous system 81, 104–109 (2000).

60. Babiec, W.E. & O’Dell, T.J. Novel Ca(2+)-dependent mechanisms regulate spontaneous release at excitatory synapses onto CA1 pyramidal cells. Journal of neurophysiology 119, 597–607 (2018).

61. Wang, Z., Low, P.A. & Vernino, S. Antibody-mediated impairment and homeostatic plasticity of autonomic ganglionic synaptic transmission. Experimental neurology 222, 114–119 (2010).

62. Grasskamp, A.T., et al. Spontaneous neurotransmission at evocable synapses predicts their responsiveness to action potentials. Front Cell Neurosci 17, 1129417 (2023).

63. Liestol, K., Maehlen, J. & Nja, A. Two types of synaptic selectivity and their interrelation during sprouting in the guinea-pig superior cervical ganglion. The Journal of physiology 384, 233–245 (1987).

64. Murray, J.G. & Thompson, J.W. The occurrence and function of collateral sprouting in the sympathetic nervous system of the cat. The Journal of physiology 135, 133–162 (1957).

65. Duan, J., Kahms, M., Steinhoff, A. & Klingauf, J. Spontaneous and evoked synaptic vesicle release arises from a single releasable pool. Cell Rep 43, 114461 (2024).

66. Ralowicz, A.J., Hokeness, S. & Hoppa, M.B. Frequency of Spontaneous Neurotransmission at Individual Boutons Corresponds to the Size of the Readily Releasable Pool of Vesicles. J Neurosci 44(2024).

67. Lichtman, J.W., Purves, D. & Yip, J.W. Innervation of sympathetic neurones in the guinea-pig thoracic chain. J Physiol 298, 285–299 (1980).

