## Supplemental Materials for "Plasticity in thoracic paravertebral sympathetic postganglionic neurons after high spinal cord transection"

**Supplementary Methods.**

Detailed equations for tSPN single compartment model:

The membrane voltage's (V) dependence on various ion currents is given by the following differential equation:

$$C_m * \frac{dV}{dt} = -I_{Na} - I_K - I_M - I_{Ca} - I_{KCa} - I_A - I_h - I_{leak} + I_{pump} + I_{ext}$$

where $C_m$ is the membrane capacitance and *I*'s refer to various ion/pump currents. $I_{ext}$ could refer either to the externally injected current under CC protocol, or the measured current under VC protocol given by

$$I_{ext} = g_E * (V_E - V)$$

where $V_E$ is the voltage at the electrode and $g_E$ is the series conductance between the electrode and the membrane.

The equations and parameters for the individual ionic/pump currents are described below.

Here ***G's*** refer to the maximal channel conductances, ***g's*** are the instantaneous conductances, ***E's*** are the Nernst equilibria and ***m,h,s,n*** etc. are the gating variables. Each gating variable of the form ***x*** is updated according to the differential equation given by:

$$\frac{dx}{dt} = \frac{x_\infty - x}{\tau_x}$$

where $\boldsymbol{\tau_x}$ describes the activation or inactivation time constant, and $\boldsymbol{x_\infty}$ describes the steady state value for ***x***.

<u>Fast TTX sensitive $Na^+$ current:</u>

$$I_{Na} = G_{Na} * m^3 * h * s * (V - E_{Na})$$

$$m_\infty = \frac{1}{1 + \exp\left(\frac{V + 41.35 - \Delta V_{Na}}{-4.75}\right)}$$

$$\Delta V_{Na} = V_{NaShift}$$

$$\tau_m = 0.3 * \exp(0.0002016125 * (V + 60.35)^2) + 0.3$$

$$\frac{dm}{dt} = \frac{m_\infty - m}{\tau_m}$$

$$h_\infty = \frac{1}{1 + \exp\left(\frac{V + 62}{4.5}\right)}$$

$$\tau_h = 23.4 * \exp(-(0.0295)^2 * (V + 75)^2) + 0.4$$

$$s_\infty = \frac{1}{1 + \exp\left(\frac{V + 40}{1.5}\right)}$$

$$\tau_s = \frac{25}{1 + \exp(\frac{V - 20}{4.5})} + 0.01$$

High-threshold long lasting $Ca^{2+}$ current:

$$I_{CaL} = G_{CaL} * \mathrm{m}_{\mathrm{CaL}} * (0.55 * h_{CaL} + 0.45 * h_{CaL2} \ \ * (V - E_{Ca})$$

$$m_{CaL\infty} = \frac{1}{1 + \exp(\frac{V + 20}{-4.5})}$$

$$\tau_{\mathrm{m}_{\mathrm{CaL}}} = 3.25 * \exp(-(0.042)^2 * (V + 31)^2) + 0.395$$

$$h_{CaL\infty} = \frac{1}{1 + \exp(\frac{V + 20}{25})}$$

$$\tau_{\mathrm{h}_{\mathrm{CaL}}} = 33.5 * \exp(-(0.0395)^2 * (V + 30)^2) + 5$$

$$h_{CaL2\,\infty} = \frac{0.2}{1 + \exp\left(\frac{V + 5}{-10}\right)} + \frac{1}{1 + \exp\left(\frac{V + 40}{10}\right)}$$

A-type $K^+$ current:

$$I_{KCa} = G_A * m_A^3 * h_A * (V - E_K)$$

$$m_{A\infty} = \frac{1}{1 + \exp(\frac{V + 38}{-18})}$$

$$\tau_{m_A} = \exp(-(0.022)^2 * (V + 75)^2) + 1.1$$

$$h_{A\infty} = \frac{1}{1 + \exp(\frac{V + 68}{7})}$$

$$\tau_{h_A} = 12 * \exp(-(0.035)^2 * (V + 40)^2) + 9$$

h-current :

$$I_h = G_h * m_h * (V - E_h)$$

$$m_{h\infty} = \frac{1}{1 + \exp\left(\frac{V + 87.6}{11.7}\right)}$$

$$\tau_{m_h activ} = 53.5 + 67.7 * \exp(\frac{V + 120}{-22.4})$$

$$\tau_{m_h deactiv} = 40.9 - 0.45 * V$$

$$\tau_{m_h} = \tau_{m_h active} \; if \; m_{h\infty} > m_{h\infty \, \mathrm{prev}} \, , else \; \tau_{m_h deactiv}$$

Leak current:

$$I_{leak} = G_{leak} * (V - E_{leak})$$

Pump current:

$$I_{pump} = J_p$$

$$\mathrm{J_p} = \mathrm{P} * \left(\frac{[\mathrm{Na_i^+}]}{[\mathrm{Na_o^+}]}\right)^3$$

where the parameter P controls the change in ion concentrations due to pump

Intracellular ion concentrations:

$$\frac{\mathrm{d}[\mathrm{Na}_i^+]}{dt} = (-A_m/F) * \{(g_{Na} + 0.21875 * G_{leak}) * (V - E_{Na}) + 3 * J_p\}$$

$$\frac{\mathrm{d}[\mathrm{K}_i^+]}{dt} = (-A_m/F) * \{(g_K + g_A + g_M + g_{KCa} + 0.78125 * G_{leak}) * (V - E_K) - 2 * J_p\}$$

where the parameter $A_m$ controls the change in membrane ion concentrations due to ion channels.

$$\frac{d[Ca_i^{2+}]}{dt} = \lambda * (-\alpha_{\mathrm{Ca}} * \mathrm{I_{CaL}} - k_{CaS}[Ca^{2+}])$$

where $\lambda$=0.01 is the ratio of free to bound [$Ca_i^{2+}$], $\alpha = 0.002$ uM/pA is the conversion factor from current to concentration, $\mathrm{k_{CaS}} = 0.024$ /ms is the [$Ca_i^{2+}$] removal rate and SCa=1 $\mu$M is the half-saturation of [$Ca_i^{2+}$].

Nernst equilibria and other parameters:

$$\mathrm{E_{Na}} = \frac{\mathrm{R * T}}{\mathrm{F}} * \log\{\frac{[Na_o^+]}{[Na_i^+]}\}$$

$$\mathrm{E_K} = \frac{\mathrm{R * T}}{\mathrm{F}} * \log\{\frac{[K_o^+]}{[K_i^+]}\}$$

$$\mathrm{E_{leak}} = \frac{0.28 * E_{Na} + E_K}{1.28}$$

$$\mathrm{E_{Ca}} = \frac{\mathrm{R * T}}{2 * \mathrm{F}} * \log\{\frac{[Ca_o^{2+}]}{[Ca_i^{2+}]}\}$$

$$\mathrm{E_h} = -31.6 \, mV$$

$$[\mathrm{Na_o^+}] = 145 \, mM$$

$$[\mathrm{K_o^+}] = 4.1\ mM$$

$$[\mathrm{Ca_o^{2+}}] = 2\ mM$$

$\mathrm{R} = 8314\,\frac{\mathrm{J}}{KgMK}$ is the ideal gas constant

$\mathrm{F} = 96500\,\frac{C}{M}$ is the Faraday constant

<u>Maximal conductances and other parameters:</u>

The model parameters for recruitment studies (**Fig 3C, Fig 8D**) are given below.

$$\{\mathrm{g_E}, \mathrm{G_{Na}}, \mathrm{G_K}, \mathrm{G_{CaL}}, \mathrm{G_M}, \mathrm{G_{KCa}}, \mathrm{G_A}, \mathrm{G_h}, \mathrm{G_{leak}}, \mathrm{C_m}, \Delta V_{Na}, \mathrm{g_{Electrode}}, \mathrm{C_{Electrode}}, \mathrm{T}, \mathrm{Pump}, \mathrm{Pump}\ A_m, E_{LeakMultiplier}\}$$

*For a chronic cell tuned across modalities:*

$$= \begin{bmatrix} 34, 11954250, 82.8, 8.0, 2, 3, 43.7, 0.23, 0.115, \\ 81, 26.3, 13.44, 0.7, 296, 0, 0.0, 0.7 \end{bmatrix}$$

*For a sham-like cell:*

$$= [\begin{matrix} 34, 2 * 11954250, 1.25 * 82.8, 8.0, 2 * 2, 3, 1.5 * 43.7, 1.5 * 0.23, 1.5 * 0.115, \\ 92, 26.3, 13.44, 0.7, 296, 0, 0.0, 0.7 \end{matrix}]$$

The model parameters for recruitment studies *f*-I curves (**Fig 7A$_4$-B$_4$**) are given below.

$$\{\mathrm{g_E}, \mathrm{G_{Na}}, \mathrm{G_K}, \mathrm{G_{CaL}}, \mathrm{G_M}, \mathrm{G_{KCa}}, \mathrm{G_A}, \mathrm{G_h}, \mathrm{G_{leak}}, \mathrm{C_m}, \Delta V_{Na}, \mathrm{g_{Electrode}}, \mathrm{C_{Electrode}}, \mathrm{T}, \mathrm{Pump}, \mathrm{Pump}\ A_m\}$$

*For a chronic cell tuned across modalities:*

$$= \begin{bmatrix} 34, 45000000, 180, 6, 0.5, 2, 190, 1, 0.5, \\ 110, 26.3, 13.44, 0.7, 296, 40000, 0.4. \end{bmatrix}$$

*For a sham-like cell:*

$$= [\begin{matrix} 34, 3 * 45000000, 2.5 * 180, 6, 0.5, 2, 1.6 * 190, 1, 3 * 0.5, \\ 1.3 * 110, 26.3, 13.44, 0.7, 296, 40000, 0.4. \end{matrix}$$

### Sup Fig 1. BP measurement

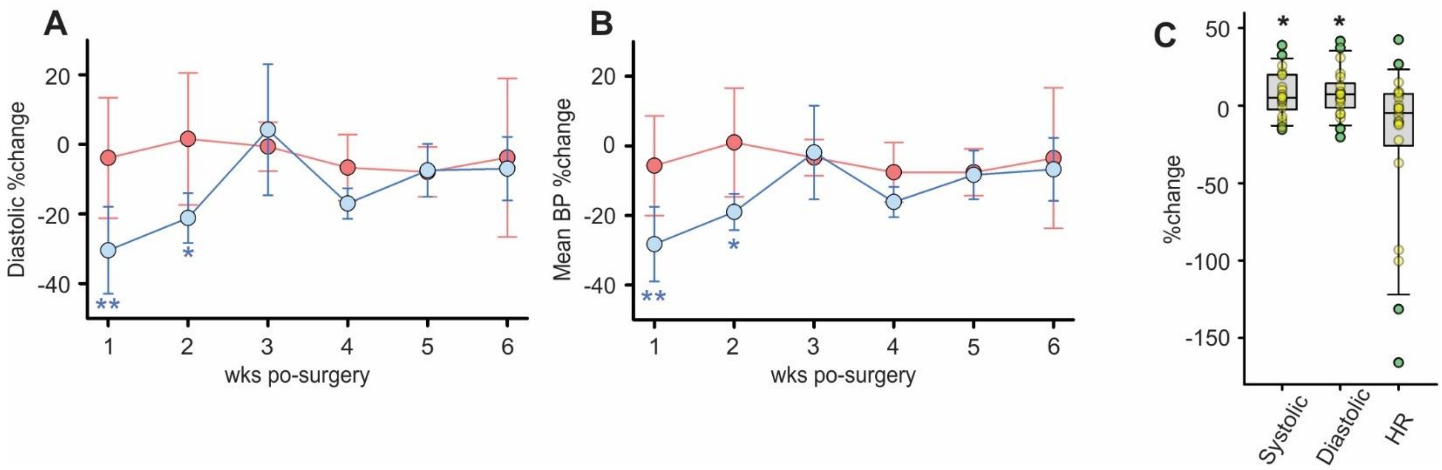


**Sup Fig 1.** Blood pressure (BP) measurements. ***A,*** Temporal change of diastolic pressure in the same recording series as in Fig 1B. Diastolic pressure was normalized with pre-surgery pressure. SCI mice (blue trace) had a significantly reduced diastolic pressure in the first two weeks after SCI (P = 0.008 at 1wk and P = 0.022 at 2wk). This reduction was not seen after sham surgery (red trace, P = 0.824 at 1wk and P = 0.874 at 2wk). ***B***, Temporal change of mean blood pressure in the same recording series. The same pattern of reduction of mean blood pressure in the first two weeks after SCI was observed. P = 0.003 at 1wk and P = 0.011 at 2wk post SCI, and P = 0.741 at 1wk and P = 0.959 at 2wk after sham surgery. * indicates significance compared to baseline. Unpaired student t-test. ***C***, Systolic pressure, diastolic pressure, and heart rate (HR) measured with full bladder was compared to those with empty bladder at 3-6 weeks post-SCI in a separate cohort (n = 22). Systolic pressure and diastolic pressure were both significantly higher with full bladder (P = 0.033 and P = 0.024, respectively), and HR was numerically lower (P = 0.189). * indicates significance between measurement with full bladder and empty bladder, paired t-test. * P < 0.05, ** P < 0.01.

**Supplementary Table 1. Selected correlations between whole population tSPN variables.**

| | | Sham | | | | SCI | | | |
|---|---|---|---|---|---|---|---|---|---|
| | | r | $R^2$ | n | P | r | $R^2$ | n | P |
| $R_{in}$ | $\tau_m$ | 0.86 | 0.73 | 17 | $1.14 \times 10^{-5}$* | 0.76 | 0.57 | 36 | $9.59 \times 10^{-8}$* |
| $R_{in}$ | $C_m$ | -0.40 | 0.16 | 17 | 0.108 | -0.24 | 0.06 | 36 | 0.161 |
| $\tau_m$ | $C_m$ | -0.02 | $2.4 \times 10^{-4}$ | 17 | 0.953 | 0.31 | 0.09 | 36 | 0.067 |
| $I_{rheo}$ | $G_{in}$ | 0.92 | 0.85 | 18 | $5.75 \times 10^{-8}$* | 0.90 | 0.82 | 35 | $9.35 \times 10^{-14}$* |
| $I_{rheo}$ | $\tau_m^{-1}$ | 0.63 | 0.40 | 17 | $6.59 \times 10^{-3}$ | 0.74 | 0.55 | 35 | $4.06 \times 10^{-7}$* |
| $I_{rheo}$ | $C_m$ | 0.49 | 0.24 | 17 | 0.046 | 0.18 | 0.03 | 35 | 0.289 |
| $I_{rheo}$ | Absolute T | 0.25 | 0.06 | 18 | 0.324 | -0.03 | $7.8 \times 10^{-4}$ | 34 | 0.876 |
| $I_{Na}$ | $I_A$ | 0.86 | 0.74 | 15 | $3.85 \times 10^{-5}$* | 0.67 | 0.45 | 32 | $3.12 \times 10^{-5}$* |
| AP Amp | $I_{Na}$ Amp | 0.83 | 0.69 | 15 | $1.34 \times 10^{-4}$* | 0.65 | 0.42 | 31 | $7.74 \times 10^{-5}$* |
| AP Amp | $I_{Na}$ Amp/ $C_m$ | 0.47 | 0.22 | 14 | 0.090 | 0.35 | 0.13 | 31 | 0.051 |
| AP Amp | $I_A$ Amp | 0.85 | 0.72 | 18 | $9.77 \times 10^{-6}$* | 0.62 | 0.38 | 34 | $1.07 \times 10^{-4}$* |
| AP Amp | $I_A$ Amp/ $C_m$ | 0.61 | 0.37 | 16 | 0.012 | 0.44 | 0.19 | 34 | $9.11 \times 10^{-3}$ |
| AP HW | $I_{Na}$ Amp | -0.61 | 0.37 | 15 | 0.016 | -0.41 | 0.16 | 31 | 0.024 |
| AP HW | $I_{Na}$ Amp/ $C_m$ | -0.30 | 0.09 | 14 | 0.295 | -0.10 | 0.01 | 31 | 0.598 |
| AP HW | $I_A$ Amp | -0.68 | 0.47 | 18 | $1.75 \times 10^{-3}$ | -0.63 | 0.40 | 34 | $5.79 \times 10^{-5}$* |
| AP HW | $I_A$ Amp/ $C_m$ | -0.43 | 0.18 | 16 | 0.100 | -0.42 | 0.17 | 34 | 0.015 |
| AP Max dV/dt | $I_{Na}$ Amp | 0.84 | 0.71 | 15 | $8.93 \times 10^{-5}$* | 0.67 | 0.44 | 31 | $4.17 \times 10^{-5}$* |
| AP Max dV/dt | $I_{Na}$ Amp/ $C_m$ | 0.38 | 0.15 | 14 | 0.178 | 0.31 | 0.10 | 31 | 0.086 |
| AP Max dV/dt | $I_A$ Amp | 0.90 | 0.81 | 18 | $3.89 \times 10^{-7}$* | 0.73 | 0.53 | 34 | $1.08 \times 10^{-6}$* |
| AP Max dV/dt | $I_A$ Amp/ $C_m$ | 0.72 | 0.51 | 16 | $1.78 \times 10^{-3}$ | 0.55 | 0.30 | 34 | $8.08 \times 10^{-4}$* |
| $f_{max}$-I slope | $R_{in}$ | 0.20 | 0.04 | 18 | 0.422 | 0.47 | 0.22 | 33 | $5.79 \times 10^{-3}$ |
| $f_{max}$-I slope | $\tau_m$ | 0.34 | 0.12 | 17 | 0.182 | 0.22 | 0.05 | 33 | 0.230 |
| $f_{max}$-I slope | $C_m$ | -0.04 | 0.002 | 17 | 0.870 | -0.25 | 0.06 | 33 | 0.163 |
| $f_{max}$-I slope | $I_{rheo}$ | -0.53 | 0.28 | 18 | 0.025 | -0.34 | 0.12 | 33 | 0.050 |
| $f_{sus}$-I slope | $R_{in}$ | -0.16 | 0.03 | 18 | 0.528 | 0.26 | 0.07 | 32 | 0.152 |
| $f_{sus}$-I slope | $\tau_m$ | -0.17 | 0.03 | 17 | 0.512 | 0.14 | 0.02 | 32 | 0.447 |
| $f_{sus}$-I slope | $C_m$ | -0.10 | 0.01 | 17 | 0.714 | -0.20 | 0.04 | 32 | 0.262 |
| $f_{sus}$-I slope | $I_{rheo}$ | -0.11 | 0.01 | 18 | 0.677 | -0.20 | 0.04 | 32 | 0.276 |
| $f_{max}$-I slope | $I_{Na}$ Amp | 0.22 | 0.05 | 15 | 0.432 | -0.14 | 0.02 | 29 | 0.466 |
| $f_{max}$-I slope | $I_A$ Amp | 0.10 | 0.01 | 18 | 0.688 | -0.34 | 0.12 | 32 | 0.057 |
| $f_{max}$-I slope | $I_K$ Amp | 0.07 | 0.004 | 16 | 0.807 | -0.37 | 0.14 | 33 | 0.035 |
| $f_{max}$-I slope | $I_{PIC}$ Amp | 0.33 | 0.11 | 13 | 0.276 | 0.36 | 0.13 | 21 | 0.113 |
| $f_{sus}$-I slope | $I_{Na}$ Amp | 0.06 | 0.004 | 15 | 0.821 | -0.22 | 0.05 | 28 | 0.252 |
| $f_{sus}$-I slope | $I_A$ Amp | -0.004 | $2.0 \times 10^{-5}$ | 18 | 0.986 | -0.37 | 0.14 | 31 | 0.041 |
| $f_{sus}$-I slope | $I_K$ Amp | 0.37 | 0.14 | 16 | 0.157 | -0.30 | 0.09 | 32 | 0.098 |
| $f_{sus}$-I slope | $I_{PIC}$ Amp | -0.0002 | $2.3 \times 10^{-8}$ | 13 | 1.000 | 0.35 | 0.12 | 21 | 0.122 |

**Supplementary Table 1.** Selective correlations between whole population tSPN parameters. **r**, Pearson's correlation coefficient; **$R^2$**, coefficient of determination; **n**, number of observations. **$R_{in}$**, input resistance; **$\tau_m$**, time constant; **$C_m$**, capacitance; **$G_{in}$**, input conductance; **I**, current; **$I_{Na}$**, sodium current; **$I_A$**, A-type potassium current; **$I_{rheo}$**, rheobase; **$I_K$**, potassium delayed outward rectifier; **$I_{PIC}$**, persistent inward current; **T**, threshold; **AP**, action potential; **Amp**, amplitude; **HW**, half width; **Max dV/dt**, maximum rising slope; **$f_{max}$**, maximum instantaneous frequency; **$f_{sus}$**, sustained firing frequency; Pearson's Product-Moment Correlation test, * in P values indicate statistically significant correlation at Šidák corrected $\alpha < 0.0014$.

**Supplementary Table 2. Summary of TH⁺ tSPNs morphology change.**

| | Naive | SCI | P value |
|---|---|---|---|
| Soma | | | |
| Max CSA (μm²) | 368.3±196.1 (61) | 376.3±215.9 (42) | 0.984 |
| Diameter (μm) | 23.2±7.0 (61) | 23.0±6.6 (42) | 0.880 |
| Dendrite | | | |
| # Primary | 3.5±2.2 (21) | 3.9±1.9 (22) | 0.531 |
| # Secondary | 5.6±2.9 (17) | 5.3±2.0 (21) | 0.652 |
| Total primary length | 177.4±142.8 (21) | 122.9±65.0 (22) | 0.112 |
| Total secondary length | 188.0±132.7 (17) | 118.2±55.2 (21) | 0.035* |
| Ave primary length | 41.6±22.2 (20) | 57.7±69.0 (21) | 0.620 |
| Ave secondary length | 35.8±32.1 (17) | 29.9±18.0 (20) | 0.867 |
| Axon | | | |
| # Primary | 1.3±0.6 (21) | 1.4±0.6 (18) | 0.245 |
| # Secondary | 2.8±1.3 (11) | 2.8±1.0 (8) | 0.962 |
| Total primary length | 145.3±133.9 (21) | 133.9±97.0 (18) | 0.933 |
| Total secondary length | 223.3±146.9 (11) | 295.8±332.7 (8) | 0.804 |

**Supplementary Table 2.** Summary of $TH^+$ tSPNs morphological properties. Data is presented as mean ± SD (number of observations). **Max CSA**, maximum cross section area; **#**, number; **Ave**, average. *$P < 0.05$, unpaired t-test is defined in method.

**Supplementary Table 3. Correlations between NPY$^+$ tSPN variables.**

| | | Sham | | | | SCI | | | |
|---|---|---|---|---|---|---|---|---|---|
| | | r | $R^2$ | n | P | r | $R^2$ | n | P |
| $R_{in}$ | $\tau_m$ | 0.84 | 0.71 | 11 | $1.11x10^{-3}$* | 0.96 | 0.92 | 12 | $8.37x10^{-7}$* |
| $R_{in}$ | $C_m$ | -0.48 | 0.23 | 11 | 0.131 | -0.29 | 0.08 | 12 | 0.369 |
| $\tau_m$ | $C_m$ | -0.11 | 0.01 | 11 | 0.756 | -0.04 | 0.002 | 12 | 0.911 |
| $I_{Na}$ | $I_A$ | 0.83 | 0.69 | 11 | $1.46x10^{-3}$ | 0.71 | 0.50 | 12 | $9.83x10^{-3}$ |
| $I_{rheo}$ | $G_{in}$ | 0.93 | 0.87 | 11 | $2.51x10^{-5}$* | 0.88 | 0.77 | 12 | $1.61x10^{-4}$* |
| $I_{rheo}$ | $\tau_m^{-1}$ | 0.66 | 0.43 | 11 | 0.028 | 0.49 | 0.24 | 12 | 0.104 |
| $I_{rheo}$ | $C_m$ | 0.55 | 0.30 | 11 | 0.080 | 0.50 | 0.25 | 12 | 0.099 |
| $I_{rheo}$ | Absolute T | 0.23 | 0.05 | 11 | 0.495 | 0.58 | 0.34 | 12 | 0.048 |
| AP Amp | $I_{Na}$ Amp | 0.83 | 0.69 | 11 | $1.45x10^{-3}$ | 0.50 | 0.25 | 12 | 0.099 |
| AP Amp | $I_{Na}$ Amp/ $C_m$ | 0.24 | 0.06 | 10 | 0.503 | 0.44 | 0.19 | 12 | 0.156 |
| AP Amp | $I_A$ Amp | 0.83 | 0.69 | 12 | $8.07x10^{-4}$* | 0.59 | 0.35 | 12 | 0.042 |
| AP Amp | $I_A$ Amp/ $C_m$ | 0.51 | 0.26 | 11 | 0.111 | 0.54 | 0.29 | 12 | 0.070 |
| AP HW | $I_{Na}$ Amp | -0.63 | 0.40 | 11 | 0.038 | -0.40 | 0.16 | 12 | 0.203 |
| AP HW | $I_{Na}$ Amp/ $C_m$ | -0.31 | 0.10 | 10 | 0.379 | -0.21 | 0.05 | 12 | 0.506 |
| AP HW | $I_A$ Amp | -0.70 | 0.49 | 12 | 0.012 | -0.72 | 0.53 | 12 | $8.31x10^{-3}$ |
| AP HW | $I_A$ Amp/ $C_m$ | -0.58 | 0.33 | 11 | 0.064 | -0.48 | 0.23 | 12 | 0.113 |
| AP Max dV/dt | $I_{Na}$ Amp | 0.82 | 0.67 | 11 | $2.05x10^{-3}$ | 0.59 | 0.35 | 12 | 0.044 |
| AP Max dV/dt | $I_{Na}$ Amp/ $C_m$ | 0.15 | 0.02 | 10 | 0.676 | 0.44 | 0.20 | 12 | 0.149 |
| AP Max dV/dt | $I_A$ Amp | 0.87 | 0.75 | 12 | $2.63x10^{-4}$* | 0.68 | 0.46 | 12 | 0.015 |
| AP Max dV/dt | $I_A$ Amp/ $C_m$ | 0.57 | 0.33 | 11 | 0.065 | 0.54 | 0.29 | 12 | 0.071 |
| $f_{max}$-I slope | $R_{in}$ | 0.22 | 0.05 | 11 | 0.519 | 0.67 | 0.45 | 11 | 0.024 |
| $f_{max}$-I slope | $\tau_m$ | 0.41 | 0.16 | 11 | 0.216 | 0.55 | 0.31 | 11 | 0.079 |
| $f_{max}$-I slope | $C_m$ | -0.12 | 0.01 | 11 | 0.737 | -0.42 | 0.18 | 11 | 0.196 |
| $f_{max}$-I slope | $I_{rheo}$ | -0.51 | 0.26 | 11 | 0.107 | -0.35 | 0.12 | 11 | 0.288 |
| $f_{sus}$-I slope | $R_{in}$ | -0.03 | $8.8x10^{-4}$ | 11 | 0.931 | 0.36 | 0.13 | 10 | 0.309 |
| $f_{sus}$-I slope | $\tau_m$ | 0.11 | 0.01 | 11 | 0.756 | 0.29 | 0.08 | 10 | 0.416 |
| $f_{sus}$-I slope | $C_m$ | 0.08 | 0.01 | 11 | 0.814 | -0.25 | 0.06 | 10 | 0.480 |
| $f_{sus}$-I slope | $I_{rheo}$ | 0.04 | $1.8x10^{-3}$ | 11 | 0.901 | -0.16 | 0.03 | 10 | 0.659 |
| $f_{max}$-I slope | $I_{Na}$ Amp | 0.23 | 0.05 | 11 | 0.496 | -0.34 | 0.12 | 11 | 0.307 |
| $f_{max}$-I slope | $I_A$ Amp | 0.23 | 0.05 | 12 | 0.482 | -0.49 | 0.24 | 11 | 0.122 |
| $f_{max}$-I slope | $I_K$ Amp | -0.09 | 0.009 | 11 | 0.785 | -0.70 | 0.49 | 11 | 0.016 |
| $f_{max}$-I slope | $I_{PIC}$ Amp | 0.54 | 0.29 | 8 | 0.166 | -0.13 | 0.02 | 6 | 0.813 |
| $f_{sus}$-I slope | $I_{Na}$ Amp | 0.46 | 0.21 | 11 | 0.151 | -0.40 | 0.16 | 10 | 0.250 |
| $f_{sus}$-I slope | $I_A$ Amp | 0.46 | 0.21 | 12 | 0.132 | -0.43 | 0.18 | 10 | 0.220 |
| $f_{sus}$-I slope | $I_K$ Amp | -0.03 | 0.001 | 11 | 0.934 | -0.46 | 0.22 | 10 | 0.177 |
| $f_{sus}$-I slope | $I_{PIC}$ Amp | 0.33 | 0.11 | 8 | 0.426 | 0.38 | 0.14 | 5 | 0.533 |

**Supplementary Table 3**. Selective correlations between NPY$^+$ tSPN parameters. **r**, Pearson's correlation coefficient; **$R^2$**, coefficient of determination; **n**, number of observations. **$R_{in}$**, input resistance; **$\tau_m$**, time constant; **$C_m$**, capacitance; **$G_{in}$**, input conductance; **AP**, action potential; **Amp**, amplitude; **HW**, half width; **Max dV/dt**, maximum rising slope; **$f_{max}$**, maximum instantaneous frequency; **$f_{sus}$**, sustained firing frequency; **I**, current; **$I_{Na}$**, sodium current; **$I_A$**, A-type potassium current; **$I_K$**, potassium delayed outward rectifier; **$I_{PIC}$**, persistent inward current; **$I_{rheo}$**, rheobase; Pearson's

Product-Moment Correlation test, * in P values indicates statistically significant correlation at Šidák corrected $\alpha < 0.0014$.

**Supplementary Table 4.** Computational model assessment of channel conductances contributing to firing properties.

| Variables | $f_{max}$-I | | $f_{sus}$-I | |
|---|---|---|---|---|
| | Slope | Shift | Slope | Shift |
| ↓ $C_m$ | ↑ | - | - | - |
| ↑ $R_{in}$ | - | ← | - | ← |
| ↓ $I_{Na}$ | - | → | - | - |
| ↓ $I_A$ | - | ← | - | ← |
| ↓ $I_K$ | ↑ | - | ↑ | - |
| ↓ $I_{CaL}$ | ↓ | - | - | ← |
| ↓ $K_{Ca}$ | - | - | - | ← |
| ↓ $I_M$ | - | - | - | - |

**Supplementary Table 4.** Computational model assessment of channel conductances contributing to firing properties. **$C_m$**, capacitance; **$R_{in}$**, input resistance; **$I_{Na}$**, sodium current; **$I_A$**, A-type potassium current; **$I_K$**, potassium delayed outward rectifier; **$I_{CaL}$,** L-type calcium current; **$I_{KCa}$**, calcium activated potassium current; **$I_M$**, M current. Note that $I_{CaL}$ activates $I_{KCa}$ in the model. ↓ $I_{CaL}$ along also leads to ↓ $I_{KCa}$ that may cause the leftward shift in $f_{sus}$-I curve.

### Sup Fig 2. PIC properties

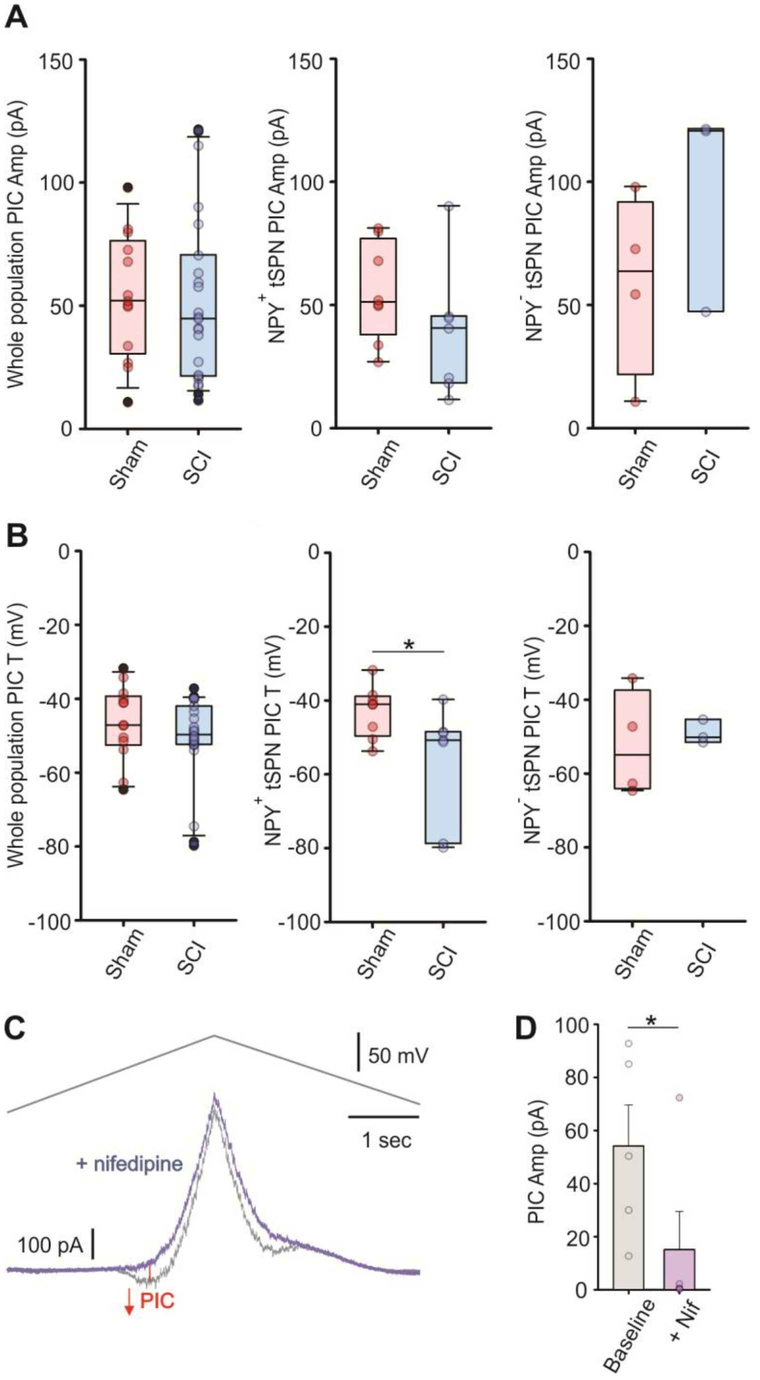


**Supplementary Figure 2.** Comparison of persistent inward current (PIC) properties in sham and SCI groups. ***A,*** Summary of PIC amplitude in the whole population (P = 0.52), NPY$^+$ tSPNs (P = 0.19), and NPY$^-$ tSPNs (P = 0.27). ***B,*** PIC threshold (T) values in the whole tSPN population were comparable in sham and SCI mice (-46.4 ± 10.1mV [n = 13] and -50.8 ± 11.8mV [n = 23]; P = 0.31). PIC threshold (T) was numerically decreased in the whole population (P = 0.307) and significantly decreased in NPY$^+$ tSPNs (-42.9 ± 7.1mV [n = 8] in sham vs -56.8 ± 15.8mV [n =7] in SCI; P = 0.04). T was unchanged in NPY$^-$ tSPNs (-52.1 ± 14.3mV [n = 4] in sham vs. -39.0 ± 3.3mV [n = 3]; P = 0.727). Unpaired t-test. ***C-D,*** Nifedipine (10 μM) completely blocked PICs in 4 out of 5 tSPNs tested (P = 0.03 comparing before and after nifedipine). One-tailed paired t-test. * P < 0.05

**Supplementary Table 5**. Correlations between sEPSC frequency and tSPNs passive membrane properties.

| | | Sham | | | | SCI | | | |
|---|---|---|---|---|---|---|---|---|---|
| Whole population | | r | $R^2$ | n | P | r | $R^2$ | n | P |
| sEPSC *f* | $C_m$ | -0.43 | 0.18 | 15 | 0.115 | -0.18 | 0.03 | 35 | 0.292 |
| sEPSC *f* | $R_{in}$ | 0.17 | 0.03 | 16 | 0.541 | 0.10 | 0.01 | 35 | 0.575 |
| sEPSC *f* | $\tau_m$ | -0.06 | 0.003 | 15 | 0.835 | -0.02 | 0.001 | 35 | 0.893 |
| $NPY^+$ | | | | | | | | | |
| sEPSC *f* | $C_m$ | -0.36 | 0.13 | 9 | 0.338 | -0.47 | 0.22 | 11 | 0.141 |
| sEPSC *f* | $R_{in}$ | 0.52 | 0.27 | 9 | 0.151 | -0.30 | 0.09 | 11 | 0.366 |
| sEPSC *f* | $\tau_m$ | 0.13 | 0.02 | 9 | 0.734 | -0.37 | 0.13 | 11 | 0.266 |

**Supplementary Table 5.** Correlations between sEPSC frequency and passive properties in whole population tSPNs and $NPY^+$ tSPNs. **r**, Pearson's correlation coefficient; $\mathbf{R^2}$, coefficient of determination; ***f***, frequency; **n**, number of observations. $\mathbf{C_m}$, capacitance; $\mathbf{R_{in}}$, input resistance; $\boldsymbol{\tau_m}$, time constant. Pearson's Product-Moment Correlation test. Šidák corrected α < 0.0085.

### Sup Fig 3. sEPSC amp and delay tau

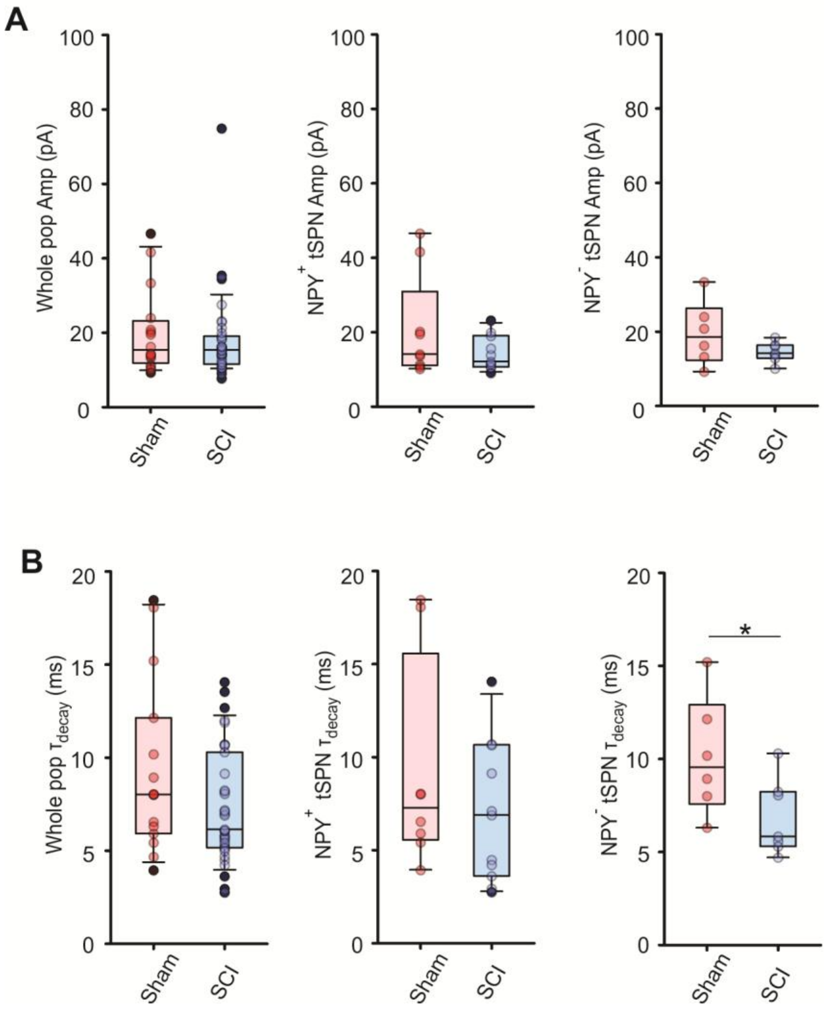


**Supplementary Figure 3.** Comparison of sympathetic preganglionic spontaneous input amplitude and decay to individual tSPNs. ***A,*** No statistical change of sEPSC amplitude in all groups after SCI ($P = 0.641$ for whole population, $P = 0.382$ for $NPY^+$ tSPNs and $P = 0.174$ for $NPY^-$ tSPNs, respectively). ***B,*** Time constant of sEPSC decay phase ($\tau_{decay}$) was numerically decreased in whole population ($P = 0.200$) and $NPY^+$ tSPNs ($P = 0.409$), but significantly decreased in $NPY^-$ population ($P = 0.046$). $^*P < 0.05$.
